# Identification of a Novel Base J Binding Protein Complex Involved in RNA Polymerase II Transcription Termination in Trypanosomes

**DOI:** 10.1101/753004

**Authors:** Rudo Kieft, Yang Zhang, Alexandre P. Marand, Jose Dagoberto Moran, Robert Bridger, Lance Wells, Robert J. Schmitz, Robert Sabatini

**Affiliations:** Department of Biochemistry and Molecular Biology, University of Georgia, Athens, Georgia, United States of America; Department of Genetics, University of Georgia, Athens, Georgia, United States of America

## Abstract

Base J, β-D-glucosyl-hydroxymethyluracil, is a modification of thymine DNA base involved in RNA Polymerase (Pol) II transcription termination in kinetoplastid protozoa. Little is understood regarding how specific thymine residues are targeted for J-modification or the mechanism of J regulated transcription termination. To identify proteins involved in J-synthesis, we expressed a tagged version of the J-glucosyltransferase (JGT) in *Leishmania tarentolae*, and identified four co-purified proteins by mass spectrometry: protein phosphatase (PP1), a homolog of Wdr82, a potential PP1 regulatory protein (PNUTS) and a protein containing a J-DNA binding domain (named JBP3). Gel shift studies indicate JBP3 is a J-DNA binding protein. Reciprocal tagging, co-IP and sucrose gradient analyses indicate PP1, JGT, JBP3, Wdr82 and PNUTS form a multimeric complex in kinetoplastids, similar to the mammalian PTW/PP1 complex involved in transcription termination via PP1 mediated dephosphorylation of Pol II. Using RNAi and analysis of Pol II termination by RNA-seq and RT-PCR, we demonstrate that ablation of PNUTS, JBP3 and Wdr82 lead to defects in Pol II termination at the 3’-end of polycistronic gene arrays in *Trypanosoma brucei*. Mutants also contain increased antisense RNA levels upstream of promoters, suggesting an additional role of the complex in regulating termination of bi-directional transcription. In addition, PNUTS loss causes derepression of silent Variant Surface Glycoprotein genes important for host immune evasion. Our results provide the first direct mechanistic link between base J and regulation of Pol II termination and suggest a novel molecular model for the role of the CTD of Pol II in terminating polycistronic transcription in trypanosomatids.

**Author Summary:** *Trypanosoma brucei* is an early-diverged parasitic protozoan that causes African sleeping sickness in humans. The genome of *T. brucei* is organized into polycistronic gene clusters that contain multiple genes that are co-transcribed from a single promoter. We have recently described the presence of a modified DNA base J and variant of histone H3 (H3.V) at transcription termination sites within gene clusters where the loss of base J and H3.V leads to read-through transcription and the expression of downstream genes. We now identify a novel stable multimeric complex containing a J binding protein (JBP3), base J glucosyltransferase (JGT), PP1 phosphatase, PP1 interactive-regulatory protein (PNUTS) and Wdr82, which we refer to as PJW/PP1. A similar complex (PTW/PP1) has been shown to be involved in Pol II termination in humans and yeast. We demonstrate that PNUTS, JBP3 and Wdr82 mutants lead to read-through transcription in *T. brucei*. Our data suggest the PJW/PP1 complex regulates termination by recruitment to termination sites via JBP3-base J interactions and dephosphorylation of specific proteins (including Pol II and termination factors) by PP1. These findings significantly expand our understanding of mechanisms underlying transcription termination in eukaryotes, including divergent organisms that utilize polycistronic transcription and novel epigenetic marks such as base J and H3.V. The studies also provide the first direct mechanistic link between J modification of DNA at termination sites and regulated Pol II termination and gene expression in kinetoplastids.

## Introduction

Termination of RNA polymerase II (Pol II) transcription of mRNAs is a tightly regulated process where the polymerase stops RNA chain elongation and dissociates from the end of the gene or transcription unit. However, the underlying termination mechanism is not fully understood. In typical eukaryotes, Pol II termination of protein-coding genes is tightly linked to the processing of the nascent transcript 3’ end (reviewed in (1)). This association ensures complete formation of stable polyadenylated mRNA products and prevents the elongating Pol II complex from interfering with transcription of downstream genes. Transcription through the polyadenylation site results in an exchange of transcription factors, resulting in the regulation of the elongation-to-termination transition, in an ordered series of events: 1) dissociation of the elongation factor Spt5, 2) Pol II pausing, 3) changes in phosphorylation status of Pol II C-terminal domain (CTD), which promotes 4) recruitment of cleavage factors and termination factors, 5) transcript cleavage and 6) termination by the 5’-3’ ‘torpedo’ exoribonuclease XRN2/Rat1.

Critical to this process is the regulation of protein phosphorylation by the major eukaryotic protein serine/threonine phosphatase, PP1. As recently demonstrated in the fission yeast *Schizosaccharomyces pombe* (*S. pombe*), the PP1 phosphatase Dis2 regulates termination by de-phosphorylating both Thr4P of the Pol II-CTD as well as Spt5 (2). This PP1-dependent dephosphorylation allows the efficient recruitment of the termination factor Seb1 as well as decreased Spt5 stimulation of Pol II elongation that enhances the capture by the torpedo exoribonuclease. Transcription termination was also affected in the *S. cerevisiae* PP1 isotype Glc7 mutant (3). PP1 action is modulated through the formation of heteromeric complexes with specific regulatory subunits (4). These regulatory protein subunits regulate PP1 by targeting the protein to specific subcellular compartments, to particular substrates, or reduce its activity towards potential substrates. The vast majority of regulators are intrinsically disordered proteins (IDPs) and bind PP1 via a primary PP1-binding motif, the RVxF motif (4–6). One of the key PP1 regulatory proteins in the nucleus is the PP1 nuclear targeting subunit (PNUTS) (7, 8). PNUTS is a multidomain protein defined by the canonical PP1 RVxF interaction motif. It also has an N-terminal domain that interacts with the DNA binding protein Tox4 and a domain near the C-terminus that interacts with Wdr82 (9). This stable multimeric complex in humans is named the PTW/PP1 complex. Targeting of the complex to chromatin is presumably due in part through associations with Tox4. While the function of Wdr82 in this complex is not known, it may mediate interactions with Pol II by recognizing Ser5-phosphorylated CTD, as it does when it is associated with the Set1 complex (10). In yeast, PP1 is associated with PNUTS and Wdr82 homologs in APT, a subcomplex of the cleavage and polyadenylation factor (11–14), and their deletion caused termination defects at Pol II-dependent genes (15, 16). In mammals, PNUTS and Wdr82 mutant cells have defects in transcription termination at the 3’ end of genes and 5’ antisense transcription at bi-directional promoters (17).

Members of the Kinetoplastida order include the human parasite *Trypanosoma brucei* that causes African Sleeping sickness. Kinetoplastids are early-diverged protozoa with unique genome arrangements where genes are organized into polycistronic transcription units (PTU) that are transcribed by Pol II. Pre-messenger RNAs (mRNA) are processed to mature mRNA with the addition of a 5’ spliced leader sequence through trans-splicing, followed by 3’ polyadenylation (reviewed in (18)). Given the close relationship between poly(A) processing and transcription termination in other eukaryotes, it is not clear how multiple functional poly(A) sites within the trypanosome PTU can be transcribed without resulting in premature termination. Therefore, a novel signal or mechanism may be involved in allowing polycistronic transcription and termination in kinetoplastids. Two chromatin factors, base J and histone H3 variant (H3.V), have recently been shown to have an impact on Pol II termination in kinetoplastids. Base J is a modified DNA base found in kinetoplastids where the glucose moiety, linked via oxygen to the thymine base, resides in the major groove of DNA (reviewed in (19)). In *T. brucei* and *Leishmania major*, J and H3.V are enriched at sites involved in Pol II termination (20–23). This includes sites within polycistronic gene clusters where (premature) termination silences downstream genes. Loss of J or H3.V leads to read-through transcription (21–26). At PTU internal termination sites this leads to increased expression of the downstream genes. This epigenetic regulation of termination is thought to allow developmentally regulated expression of specific transcriptionally silent genes. In *T. brucei*, this includes the expression of variant surface glycoprotein (VSG) genes involved in antigenic variation during bloodstream infections.

It is currently unclear how these epigenetic marks are involved in Pol II termination, since very little is understood regarding the mechanism of Pol II termination in kinetoplastids. Equally unclear is what regulates the specific localization of these marks, including base J, in the genome. Base J is synthesized in a two-step pathway in which a thymidine hydroxylase (TH), JBP1 or JBP2, hydroxylates thymidine residues at specific positions in DNA to form hydroxymethyluracil (hmU) (27), followed by the transfer of glucose to hmU by the glucosyltransferase, JGT (28, 29) (reviewed in (19, 30)). The TH enzymes, JBP1 and JBP2, contain a TH domain at the N-terminus (27, 31–34). JBP1 has a J-DNA binding domain in the C-terminal half of the protein that is able to bind synthetic J-DNA substrates *in vitro* and bind chromatin in a J-dependent manner *in vivo* (35–38). The ability of JBP1 to bind J-DNA is thought to play a role in J propagation and maintenance. JBP2 does not bind the modified base directly, but is able to bind chromatin in a base J independent manner, presumably via the C-terminal SWI2/SNF2 domain (31, 35). The JGT lacks DNA sequence specificity, and can convert hmU to base J *in vivo* regardless of where it is present (39–41). This and other analysis of J synthesis indicate that JBP1 and JBP2 are the key regulatory enzymes of J synthesis.

In this study, we identify a new J-binding protein (called JBP3) in kinetoplastids that is present in a complex containing PP1, Wdr82 and a putative orthologue of PNUTS. To characterize the role of this (PJW/PP1) complex in transcription termination, we investigated the consequence of mutants using RNAi in *T. brucei*. Ablation of JBP3, Wdr82 or PNUTS in *T. brucei* causes read-through transcription at termination sites. As we previously demonstrated following the removal of base J and H3.V, these defects include transcription read-through at termination sites within Pol II transcribed gene arrays and the silent Pol I transcribed VSG expression sites, leading to de-repression of genes involved in parasite pathogenesis. Furthermore, ablation of JBP3, Wdr82 or PNUTS results in expression of genes upstream of Pol II transcription start sites presumably representing a previously unappreciated role of termination of antisense transcription from gene promoters in trypanosomes. Overall these findings provide a first look at mechanisms involved in Pol II transcription termination in kinetoplastids and a direct link between base J and termination.

## Results

### Affinity purification and identification of JGT-associated proteins in Leishmania

In this study, we chose to identify the proteins that co-purify with JBP1, JBP2 and JGT, in order to understand the regulation of J-biosynthesis*. Leishmania tarentolae* was chosen as an experimental system, because it provides an easily grown source of high densities of parasites that synthesize base J. The proteins were cloned into the pSNSAP1 vector, carrying a C-terminal tag composed of a protein A domain, the TEV protease cleavage site, and the streptavidin binding peptide (42). Purification of the complexes was carried out as described in Materials and Methods. Separation of the final eluate from the JGT tagged cell line by SDS-PAGE and staining of the proteins by SYPRO Ruby, revealed a dominant protein band of ~120 kDa, representing JGT-S and several co-purified protein bands (Fig 1A). As a control, no specific proteins were identified from the purification from untagged wildtype cells. In contrast, and unexpectedly, no co-purified proteins were identified from the JBP1-S or JBP2-S purification either by SDS-PAGE or mass spectrometry analysis. Therefore, we continued the analysis on the JGT purification. Liquid chromatography-tandem mass spectrometry (LC-MS/MS) of the JGT-S purification revealed a total of five unique proteins not present in the negative control purification (Table 1). These proteins were required to have at least two unique peptides and a Seaquest score of at least 100. As expected, JGT-S was recovered in the eluate. However, four additional potential JGT associated proteins were discovered with (hypothetical) molecular masses of 74, 42 and 29 kDa. One of these JGT-interacting proteins (42 kDa; LtaP15.0230) was protein phosphatase 1 (PP1), which contains a PP1 catalytic domain (Fig 1B). The other 42-kDa protein (LtaP32.3990) contains three WD repeat domains and has been identified as a homologue of Wdr82/Swd2 (human/yeast). The remaining two JGT interacting proteins had not been previously characterized. We named the 74-kDa protein JBP3 (LtaP36.0380) because it has a domain with homology to the base J DNA binding domain of JBP1 (Fig 1B and S1) and we demonstrate its ability to bind J-DNA (see below). The 29-kDa JGT interacting protein was named PNUTS (LtaP33.1440) since it contains a conserved RVxF PP1 interactive domain (Fig 1C and S2 Fig) within an apparent intrinsically disordered region of the protein and is a part of a complex similar to the PTW/PP1 complex in humans (where JBP3 may represent a functional homolog of Tox4). PP1 interactive proteins such as PNUTS are highly disordered in their unbound state and fall in a group of intrinsically disordered proteins (IDPs) (4–6). This intrinsic flexibility is important for the formation of extensive interactions with PP1 (6, 43, 44). A bioinformatics analysis using the DISOPRED3 program (45, 46), which scores for the occurrence of disorder-inducing amino acids, predicts a majority of TbPNUTS is disordered (S2 Fig). Similarly, the Compositional profiler (47) shows PNUTS is enriched in major disorder-promoting residues and depleted in major order-promoting residues. This inherent disorder may explain why TbPNUTS migrates slower in SDS-PAGE than predicted (see S3A Fig). To confirm the complex, we subsequently performed tandem affinity purifications with tagged JBP3 and Wdr82. Reciprocal purification of JBP3 and Wdr82 resulted in the identification of JGT, PNUTS, PP1, JBP3 and Wdr82 with a significant number of unique peptide hits (Table 1). These data indicate that in Leishmania JGT associates with a protein complex composed of PNUTS, PP1, JBP3 and Wdr82 similar to the PTW/PP1 complex in humans, shown to be involved in transcription termination (Fig 1F). While JBP3 may be a functional homologue of the human Tox4 DNA binding protein (see below), JBP3 appears to be conserved only in the genus trypanosome. Based on this similarity, we now refer to this complex as PJW/PP1 in Leishmania.

**Fig 1.**
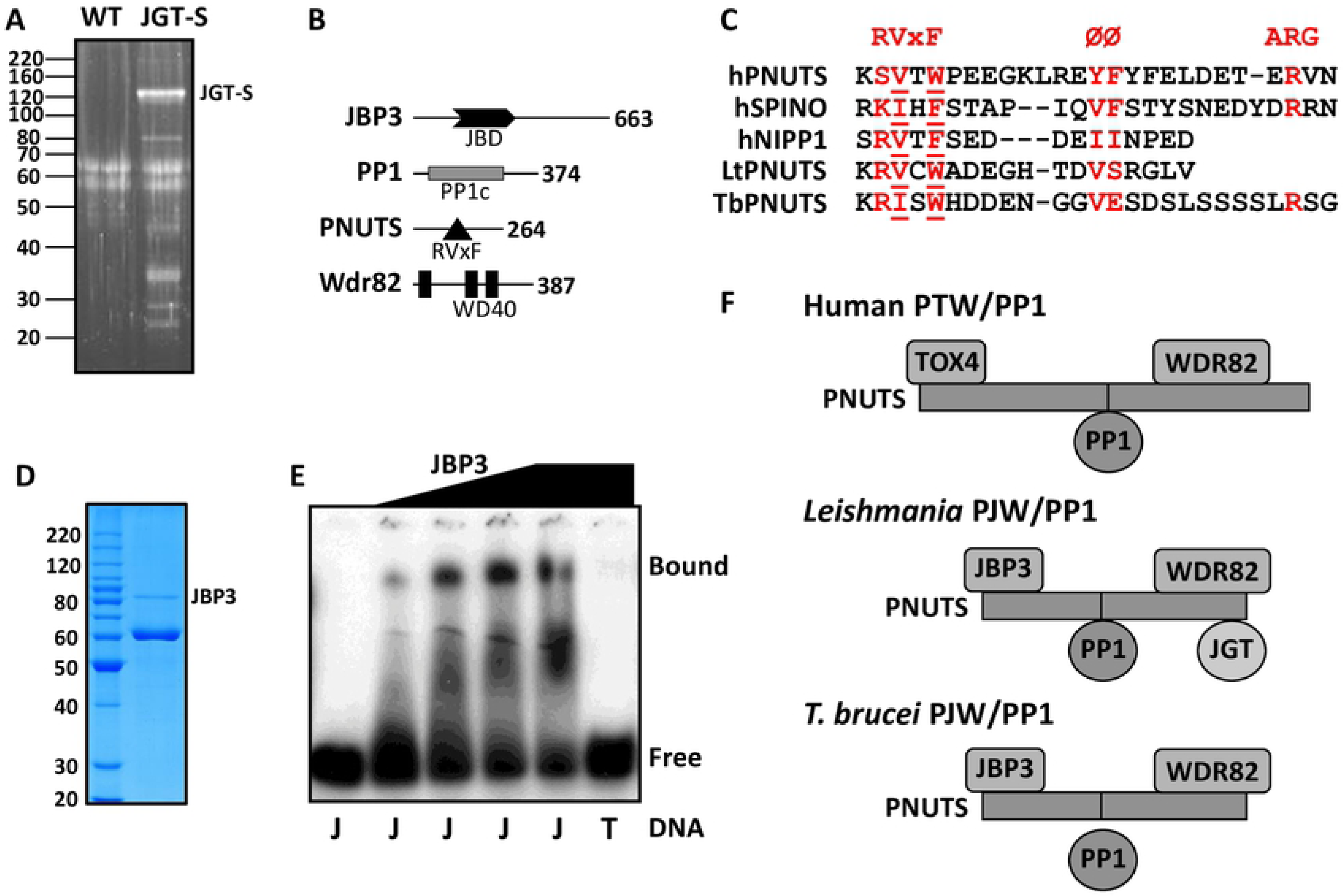
Identification of a novel phosphatase complex in *Leishmania tarentolae*. (A) Proteins recovered from tandem affinity purification from wild-type *L. tarentolae* extracts (WT) and from cells expressing the strep-tagged version of JGT, were analyzed on SDS-PAGE gel and stained with Sypro Ruby. (B) Summary of the PJW/PP1 complex. The domain structure of each component in the complex is schematically shown. JBD, J-DNA binding domain; RVxF motif, PP1 docking motif; PP1c, catalytic domain of protein phosphatase 1; WD40, WD40 repeat. The number of amino acids in each component is indicated. (C) Structure-based sequence alignment of the PNUTS, spinophilin PP1 and NIPP1 interactive domains in humans is compared with LtPNUTS (residues 95-112) and TbPNUTS (residues 139-164), where x is any amino acid and Ø represents a hydrophobic amino acid. Critical residues in the RVxF motif are underlined. (D) Purification of recombinant LtJBP3. His-tagged rJBP3 expressed in *E. coli* was purified by metal affinity and size exclusion chromatography and analyzed by SDS-PAGE/Coomassie staining. The major copurified protein is the *E. coli* molecular chaperone GroEL. The migration of protein molecular mass standards (in kDa) is shown on the left. (E) Gel shift assays for modified and unmodified DNA substrates interacting with JBP3. 0.3 pmol radiolabeled J-DNA (J) was incubated with 0, 0.2, 0.38, 0.57 and 1 pmol of JBP3 and 0.3 pmol radiolabeled unmodified DNA (T) was incubated with 1 pmol of JBP3. (F) Models for the PJW/PP1 complex in Leishmania and *T. brucei*. The models are based on the human PTW/PP1 complex where PNUTS acts as scaffold and the DNA binding protein and Wdr82 bind to the N- and C-terminus, respectively, and PP1 binds via the RVxF PP1 interaction motif (indicated by the line). PP1 is not stably associated with the complex in *T. brucei* (see discussion).

**Table 1.**
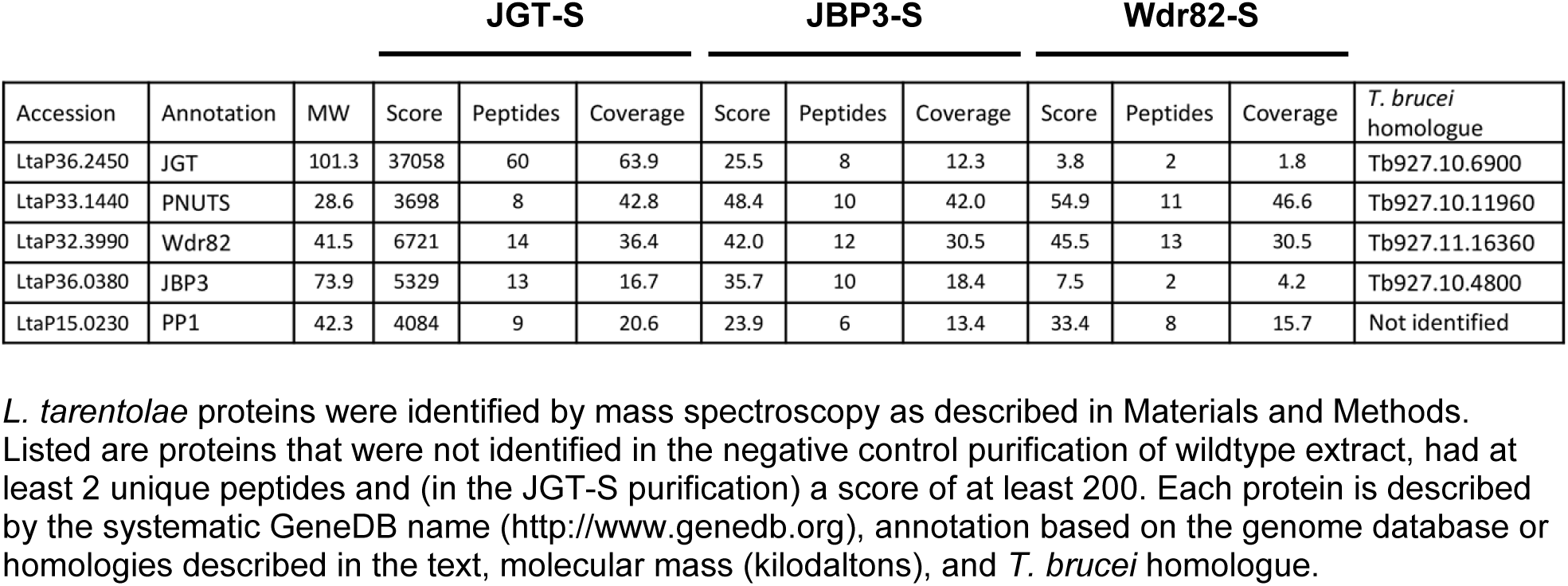
Mass spectrometric identification of JGT, JBP3 and Wdr82 purification products

### JBP3 is a J-DNA binding protein

We noticed that a region of the LtaP36.0380 protein (amino acids 101-277) has sequence similarity with the J-binding domain of JBP1 (S1A and B Fig). A structural model of the Lt JBP1 JBD based on the X-ray crystallography-based structure (PDB ID 2xseA) is shown in S1C Fig. As expected from the sequence similarities between the kinetoplastid JBD for JBP1 and the putative JBP3, the 3D models generated by the comparative modeling program I-TASSER (48, 49) were of high quality, with TM-score value of 0.727. The sequence identity and similar domain composition to the JBD of JBP1 supported our contention that the LtaP36.0380 protein is a J-binding protein, subsequently named JBP3.

We have previously developed a rapid isolation procedure for His-tagged recombinant JBP1 produced in *E. coli* and gel shift assays to characterize J-DNA binding activities (37, 38). We utilized a similar approach to characterize JBP3 *in vitro* DNA binding activity. The recombinant JBP3 obtained, as judged by its appearance in Coomassie-stained gels (Fig 1D), was suitable for investigating the specific interaction of JBP3 and J-modified DNA (J-DNA). To determine the ability of JBP3 to bind J-DNA, we used the gel shift assay to investigate the binding of JBP3 to J-DNA duplex (VSG-1J) that has a single centrally located J modification compared with the same duplex without any base J (VSG-T). The J-DNA substrate was incubated with increasing amounts of JBP3 protein, and the complex was analyzed on native gels. The results of the gel shift assay in Fig 1E, show that the amount of free J-DNA decreases with increasing concentrations of JBP3, concomitant with formation of the JBP3/J-DNA complex. In contrast, incubation of the unmodified DNA substrate with the highest concentration of JBP3 resulted in no visible complex. Thus, JBP3 is a J-DNA binding protein.

### Characterization of the putative phosphatase complex containing Wdr82, JBP3 and PNUTS in T. brucei

To study the function of the PJW/PP1 complex *in vivo*, we switched to *T. brucei* due to the benefits of forward and reverse genetics available in this system. However, while all other components are easily identified (see Tables 1 and 2), a PP1 homologue identified in the Leishmania PJW/PP1 complex is not present in the *T. brucei* genome (50, 51). To characterize the complex in *T. brucei*, and identify the TbPP1 protein component, we used a TAP tagging approach with a PTP epitope tag (52) to identify TbPNUTS-interacting proteins. We generated a clonal procyclic form (PF) *T. brucei* cell line expressing a C-terminal PTP-tagged TbPNUTS protein from its endogenous locus. This cell line was used for the TAP procedure. Briefly, a crude protein extract was first purified by IgG affinity chromatography, and the TEV protease was used to cleave off the Protein A portion of the PTP tag. Subsequently, the TEV protease eluate underwent anti-Protein C affinity purification, and the final purified products were eluted with EGTA. The concentrated proteins were then trypsin-digested and analyzed by LC-MS/MS. As a control, we analyzed the eluate of a comparable purification of extract from wild-type *T. brucei*. As expected, TbPNUTS was identified along with Wdr82 and JBP3 (Table 2). HSP70 was also identified as a potential PNUTS-interacting protein. However, JGT was not identified as part of the TbPNUTS complex along with no PP1 homologue.

**Table 2.** Mass spectrometric identification of PNUTS co-purified proteins in *T. brucei*

**PNUTS PTP**
| Accession | Annotation | MW | Score | Peptides | Coverage |
| --- | --- | --- | --- | --- | --- |
| Tb927.10.11960 | PNUTS | 35.4 | 711 | 21 | 83 |
| Tb927.11.16360 | Wdr82 | 37.9 | 275 | 12 | 52 |
| Tb927.10.4800 | JBP3 | 65.5 | 355 | 27 | 48 |
| Tb927.11.3110 | Hsp70 | 71.5 | 157 | 21 | 40 |

List of PNUTS-PTP co-purified proteins, identified by LC/MS/MS, that were not a common contaminant of previous tandem affinity purifications in *T. brucei*, were not present in the negative control purification of wildtype extract, had at least two unique peptides and a score of at least 200. Protein annotation and accession numbers are from the *T. brucei* 927 database at www.TriTrypDB.org.

Co-immunoprecipitation studies were performed to further assess the PNUTS-containing complex in *T. brucei*. Representative components of the putative PP1 complex were analyzed by western blot following immunoprecipitation to assess the authenticity of protein interactions identified by mass spectrometry. To do this, we utilized BSF *T. brucei* parasites due to the ability to efficiently tag endogenous alleles. We generated *T. brucei* cell lines expressing PTP-tagged versions of PNUTS, JBP3, Wdr82 and CPSF73 (negative control) (S3A Fig) along with HA-tagged versions of JGT, Wdr82 and JBP3 (Fig 2A and S3B Fig). PTP-PNUTS immunoprecipitation recovers HA-Wdr82 and HA-JBP3, but not the La negative control (Fig 2A). Similarly, PTP-JBP3 immunoprecipitation recovers HA-Wdr82 (S3B Fig). In contrast, no detectable HA-Wdr82 or HA-JGT is recovered in PTP-CPSF73 immunoprecipitates (Fig 2A and S3B Fig). Furthermore, JGT does not co-purify when PNUTS, JBP3 or Wdr82 is pulled down (S3B Fig), nor does Hsp70 co-purify when PNUTS or JBP3 is pulled down (S3C Fig).

**Fig 2.**
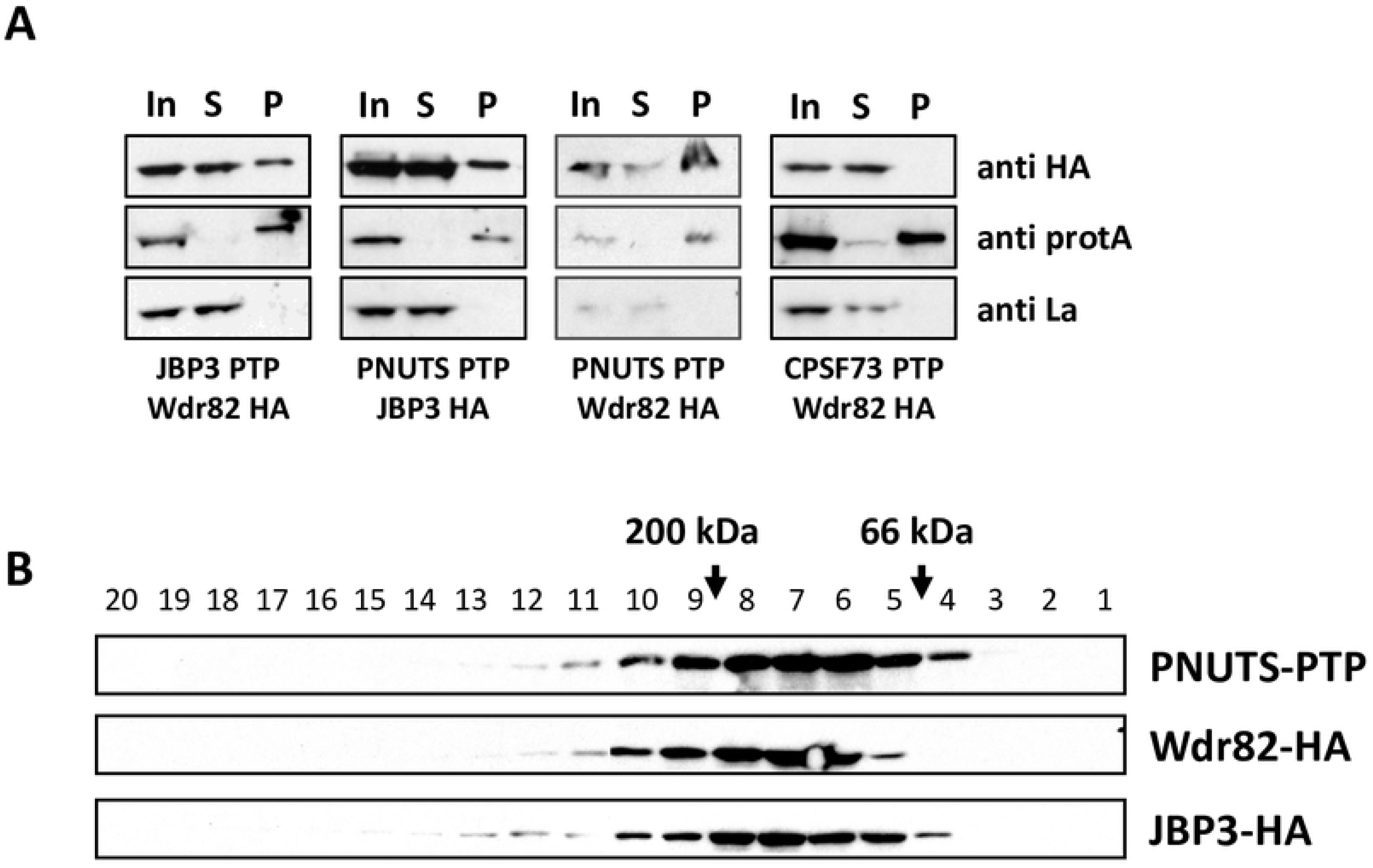
Characterization of the PJW/PP1 complex in *T. brucei*. (A) Co-immunoprecipitation of JBP3, Wdr82 and PNUTS. Cell extracts from bloodstream form *T. brucei* cells that endogenously express PTP- and HA-tagged versions of the indicated proteins were purified by protein A affinity and analyzed by western blotting with anti-HA, anti-protein A and anti-La. (B) PNUTS, Wdr82, and JBP3 co-migrate following sucrose gradient fractionation. Cell extracts from BSF *T. brucei* cells that endogenously express PNUTS-PTP and JBP3-HA/WDR82-HA were loaded onto 5-50% sucrose gradients and analyzed by density gradient centrifugation. Equal volumes of each fraction were analyzed by western blotting. Migration of PNUTS, JBP3 and WDR82 are shown in the gradient. Molecular markers were applied to a parallel gradient.

To further characterize the PNUTS complex, extracts recovered from *T. brucei* cells expressing epitope-tagged components were analyzed by western blot following sucrose fractionation. Analysis of *T. brucei* cells expressing PTP-PNUTS and HA-JBP3 or HA-Wdr82 indicate Wdr82, JBP3 and PNUTS co-migrate at <200kDa. These results indicate that JBP3, Wdr82 and PNUTS form a stable complex of <200kDa. Summation of the predicted size of the three complex components (~180 kDa) agrees with the observed mass of the PJW complex and suggests a 1:1 stoichiometry for the subunits. Taken together, data from co-immunoprecipitation, sucrose gradient analysis, and identification of PNUTS-, Wdr82-, and JBP3-associated proteins by mass spectrometry indicate that Wdr82, JBP3 and PNUTS comprise a stable PJW complex in kinetoplastids (Fig 1F). In contrast to the complex in Leishmania, the stable purified *T. brucei* complex lacks both JGT and PP1.

LtPP1 is found to be stably associated in PJW/PP1 complex, possibly through its putative PP1-interacting protein, PNUTS. PNUTS is present in both the *T. brucei* and Leishmania complex and share a conserved RVxF motif (Fig 1C). Moreover, a similar human PTW/PP1 complex suggests that PP1 confers the complex phosphatase activity critical for its regulation in Pol II termination (2, 3, 53, 54). The apparent lack of PP1 in the *T. brucei* complex is therefore surprising. To identify the TbPP1 component, we directly tested two of the eight PP1 genes in *T. brucei* for interactions with the complex by co-IP. PP1-1 has the highest sequence homology to the PP1 involved in termination in yeast and humans (51). PP1-7 has been identified in the nucleus of *T. brucei* (55). However, neither of these PP1 proteins associate with TbPNUTS in the co-IP (S3C Fig). Alternatively, PP1 is not stably associated with the complex in *T. brucei*. This is consistent with one of three replicates of LtJGT purification that resulted in all components of the complex except PP1 (S1 Table), suggesting PP1 is the least stable component of the Leishmania complex. It has been demonstrated that the phosphorylation of Ser residue(s) within or close to the RVxF motif of PP1 regulatory subunits, including PNUTS, disrupts the binding of this motif to PP1 (56–59). Treatment of *T. brucei* extract with calf intestinal phosphatase results in a shift of TbPNUTS mobility on SDS-PAGE (S3D Fig), suggesting TbPNUTS is phosphorylated *in vivo*. These data suggest PP1 is an (unstable) component of the PJW/PP1 complex in *T. brucei* where association may be regulated by PNUTS phosphorylation.

### Downregulation of PNUTS, Wdr82 and JBP3 causes defects in RNA Pol II transcription termination at the 3’end of PTUs

If PNUTS, Wdr82 and JBP3 interact with each other, one would predict that RNAi against the individual components would give the same phenotype. We therefore analyzed the role of the PJW/PP1 complex in transcription termination in BSF *T. brucei*. We have previously shown that base J and H3.V are present at termination sites within a PTU where loss of either epigenetic mark results in read-through transcription and increased expression of downstream genes (21, 24, 25). Because Pol II elongation and gene expression is inhibited prior to the end of these PTUs, we refer to this as PTU internal termination. For example, base J and H3.V are involved in terminating transcription upstream of the last two genes (Tb927.5.3990 and Tb927.5.4000) in a PTU on chromosome 5 (Fig 3, top) where deletion of H3.V or inhibition of base J synthesis leads to read-through transcription (21, 24). The resulting read-through RNAs become observable in a manner similar to those seen in other systems when termination mechanisms become incapacitated by experimental manipulation. The presence of an open reading frame downstream of the termination site allows an additional measure of read-through where nascent RNA is processed to stable capped and polyadenylated mRNA species. As such, the loss of either epigenetic mark in *T. brucei* leads to generation of nascent RNA extending beyond the termination site and expression of the two downstream genes (21, 24). To investigate the physiologic function of the PJW/PP1 complex, we analyzed inducible RNAi ablation of Wdr82, JBP3, and PNUTS in BSF *T. brucei*. As shown in Fig 3, induction of RNAi against Wdr82, JBP3 and PNUTS and ablation of mRNA levels from 30-60% leads to reduced parasite growth, indicating the essential nature of these genes and thus the PJW/PP1 complex in BSF *T. brucei*. We also detect evidence of read-through transcription at the representative PTU internal termination site on chromosome 5 upon ablation of the three factors. RT-PCR using oligos flanking the termination site (see diagram, Fig 3A) detects increased RNA upon ablation of PNUTS, Wdr82 and JBP3 (Fig 3C). As a control a separate RT-PCR utilized the same 5’ primer and a 3’ primer immediately upstream of the termination site. We have previously shown that an RNA species spanning the termination site is indicative of read-through transcription and is only detected following the loss of base J or H3.V, due to continued transcription elongation at termination sites (21, 24, 25). Consistent with read-through transcription, both genes downstream of the termination site are significantly de-repressed upon the ablation of the three components of the PJW/PP1 complex, in contrast to genes upstream (Fig 3D). In contrast, no significant termination defects are detected upon ablation of a negative control, acidocalcisome VA a protein (Fig 3 C and D). VA a has been shown to be an essential gene in *T. brucei* and ablation results in similar growth defects as seen in PJW/PP1 complex (60) (Fig 3B). Therefore, the read-through defects measured in the PNUTS, Wdr82 and JBP3 mutants are presumably not the result of indirect effects of dying cells. These results suggest that PNUTS, Wdr82 and JBP3, possibly functioning in the PJW/PP1 complex, have a role in regulating Pol II termination.

**Fig 3.**
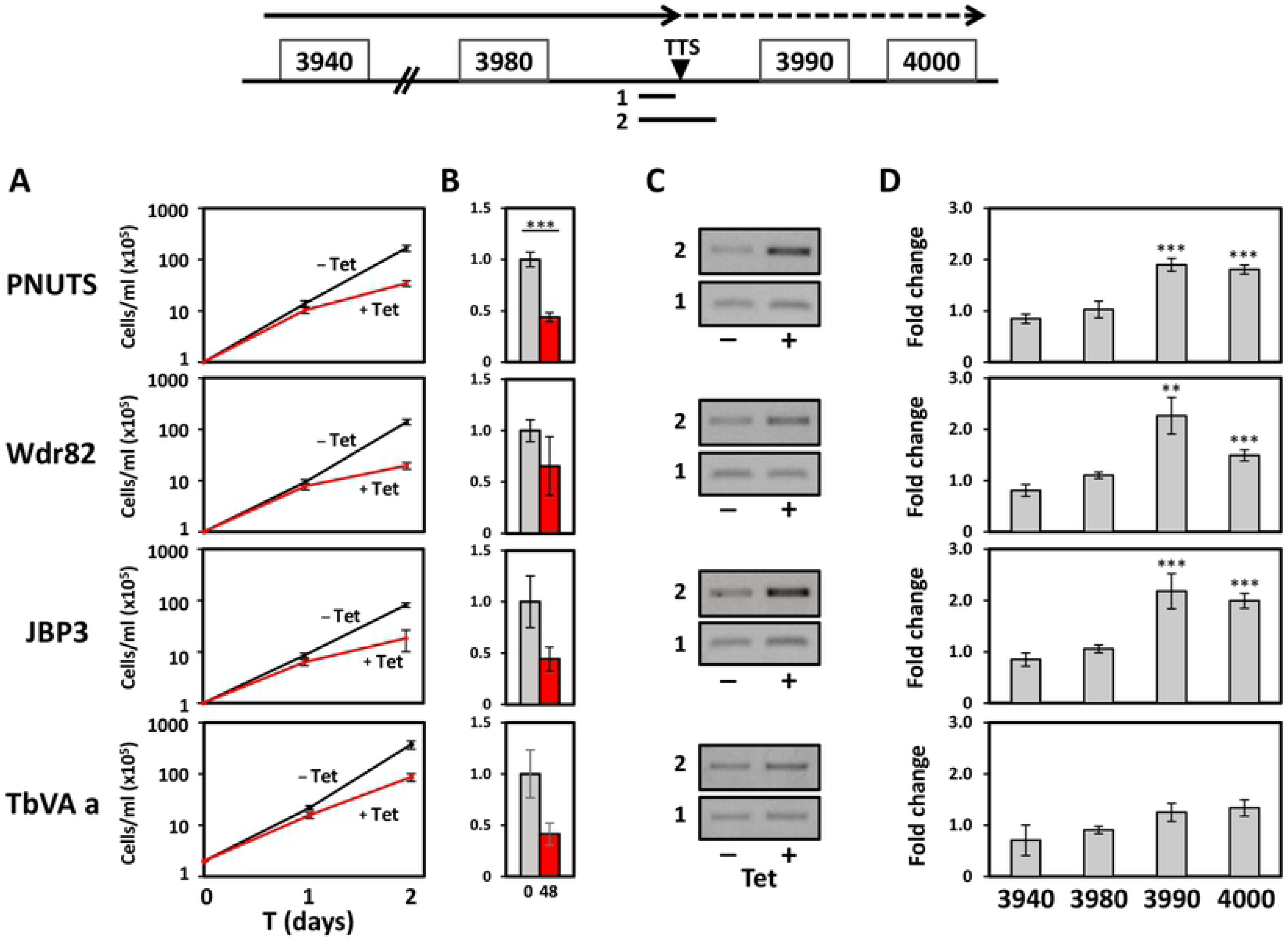
PNUTS, Wdr82 and JBP3 are involved in Pol II transcription termination in *T. brucei*. (A) PNUTS, JBP3 and Wdr82 are essential for cell viability of the infectious BSF *T. brucei*. Cell growth was arrested upon PNUTS, JBP3 and Wdr82 mRNA ablation by RNAi. VA a; acidocalcisome VA a protein. (B) Depletion of mRNA upon Tet induction of RNAi by qRT-PCR analysis. P values were calculated using Student’s t test. ***, p value ≤ 0.001. (C and D) Analysis of Pol II termination. Above; Schematic representation of a termination site on chromosome 5 where Pol II has been shown to terminate prior to the last two genes (Tb927.5.3990 and Tb927.5.4000) in the PTU. The dashed arrow indicates readthrough transcription past the TTS that is regulated by base J and H3.V. Solid lines below indicate regions (1 and 2) analyzed by RT-PCR. (C) RT-PCR analysis of nascent RNA. cDNA was synthesized using random hexamers and PCR was performed using the appropriate reverse primer for each region (1 and 2) plus an identical forward primer. (D) RT-qPCR analysis of genes numbered according to the ORF map above. Transcripts were normalized against Asf1, and are plotted as the average and standard deviation of three replicates. P values were calculated using Student’s t test. **, p value ≤ 0.01; ***, p value ≤ 0.001.

To further explore the role of the PJW/PP1 complex in the regulation of termination and whether the complex functions similarly across the genome, we performed stranded mRNA-seq to compare the expression profiles of PNUTS RNAi cells with and without tetracyclin induction. This led to the detection of 709 mRNAs that are increased at least 3-fold upon ablation of PNUTS (P_adj_<0.05) (S4A Fig and S2 Table). In contrast, no mRNAs are downregulated. Of the 3-fold upregulated genes, a majority represent VSGs, ESAGs and Retrotransposon Hot Spot proteins (RHS) that are repressed in bloodstream *T. brucei* and are localized at the end of Pol II transcribed PTUs located within the chromosomes or in subtelomeres. The location of the genes with >3-fold upregulation was mapped (Fig 4A and S5). Interestingly, these genes were closely located at regions flanking PTUs or subtelomeric regions, suggesting a role of PNUTS in genome-wide transcription, specifically at transcription initiation and termination sites, as well as subtelomeres. The PTU flanking regions represent transcription termination sites (TTS) and transcription start sites (TSS) within the so-called convergent strand switch region (cSSR) and divergent strand switch region (dSSR), respectively. We have previously shown that base J is localized within the cSSRs and dSSRs flanking PTUs and involved in Pol II termination. Therefore, we wanted to first confirm the association with J at termination sites with changes in gene expression we observed in the PNUTS mutant. If we analyze genes within 1kb of base J genome-wide, we see a correlation with upregulation of gene expression versus a similar analysis of the same number of randomly selected genes across the genome (S4B Fig). In contrast, the 4 genes upregulated in the VA a mutant lack any association with base J. The upregulated genes in the PNUTS mutant located at the 3’ end of PTUs, including the VSG gene (Tb927.5.3990) analyzed in Fig 3, map to regions downstream of base J and H3.V. The RNA-seq results confirm our initial RT-PCR analysis of nascent and steady-state RNA indicating a role of PNUTS, Wdr82 and JBP3 regulating termination and expression of downstream genes (Fig 3). Therefore, consistent with our analysis of the J/H3.V mutants, in the PNUTS mutant we observe similar increases in the expression of genes downstream of PTU internal termination sites, which we previously demonstrated is caused by a defect in Pol II transcription termination resulting in readthrough transcription (21, 24, 25).

**Fig 4.**
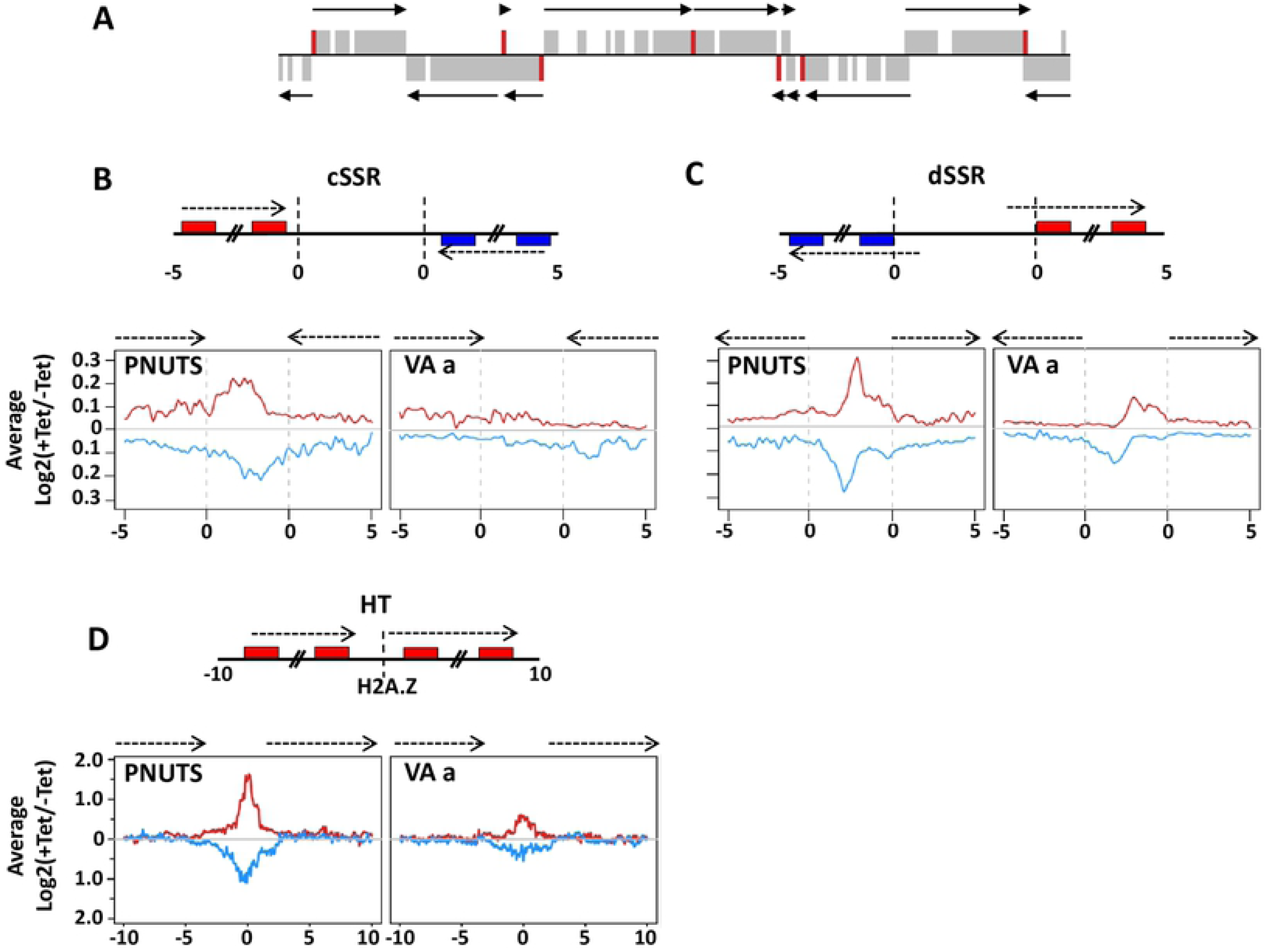
TbPNUTS affects transcription termination at 3’ end and 5’ end of PTUs. Levels of transcripts downstream of TTSs and upstream of TSSs increased upon TbPNUTS ablation. (A) A core section of chromosome 10 is shown; genes that were upregulated >3-fold upon ablation of PNUTS are highlighted in red. Arrows indicate the direction of transcription of PTUs. (B and C) Diagram of a cSSR (B) and dSSR (C): boxes are genes in the PTUs and arrows indicate PTUs and direction of transcription. In cSSR, the poly A sites of the final gene in the PTU (indicated by the transcriptome) are marked by dotted line (0 kb). In dSSR, the dotted line (0 kb) indicates the 5’ end of the first gene in the PTU (according to the transcriptome). Thus, the TSS is located further upstream, within the dSSR. Numbers refer to distance from SSR (kb). Below: Metagene profile of total sense and antisense RNA-seq signal over the SSRs and 5kb upstream and downstream regions into the PTUs. Fold changes comparing transcription levels between day 0 and day 2 following induction of RNAi were calculated and plotted over the indicated regions genome-wide. Red and blue lines indicate RNAs from the top and bottom strand, respectively, as indicated on the diagram above. PNUTS, PNUTS RNAi; VA a, RNAi VA a. (D) Above: Diagram of a HT site. Boxes are the genes and arrows indicate direction of transcription as in C. The center of the H2A.Z peak is marked by a dotted line. Numbers refer to distance from center of H2A.Z peak (kb). Below: Metagene profile of the fold changes in RNA-seq reads over the HT sites and 10 kb upstream and downstream of the H2A.Z peak, as described in C.

A remaining question was whether PNUTS also regulated termination at the 3’-end of gene clusters resulting in transcription into the SSR. Our previous RNA-seq analyses in *T. brucei* indicated that H3.V/J regulated termination at the 3’-end of PTUs, where mutants resulted in Pol II elongation into the dual strand transcription regions at cSSRs (21, 24). To visualize the differences between WT and TbPNUTS mutant, fold changes between –Tet and +Tet induction of RNAi were plotted (S6 Fig). While sense transcription remained largely unaffected throughout all 11 megabase chromosomes following the loss of PNUTS, significant fold increases of antisense transcription were observed near transcription borders of PTUs, including downstream of normal transcription termination sites at the 3’-end. To take a closer look at the increased level of transcripts at these sites, and determine whether they are due to transcriptional readthrough, forward and reverse reads mapping to 5kb flanking and within cSSRs were counted and RPKM values were generated. As shown in Fig 4B, cSSRs are computationally defined regions where coding strands switch based on the transcriptome. To see the difference between WT and PNUTS mutant, changes in transcript abundance upon PNUTS and VA a ablation were plotted. A metaplot summarizing the readthrough defect for all cSSRs is shown in Figure 4B. We observed an increase in RNA extending into the cSSR in the PNUTS mutant, and no change in the VA a mutant control. Boxplots comparing the median RPKM values for SSRs also indicated that cSSRs were upregulated in the PNUTS mutant and the differences were statistically significant (S7 Fig). All together these results indicate that the loss of PNUTS, JBP3, and Wdr82 result in defects in transcription termination at the 3’-end of PTUs as we described in the H3.V/J mutant.

### PNUTS, Wdr82 and JBP3 are involved in Pol II transcription termination at 5’ end of PTUs

Based on the initial RT-PCR analysis of the internal termination site in PNUTS, JBP3 and Wdr82 knockdowns (Fig 3), we expected RNA-seq to illustrate defects in termination at 3’-end of PTUs genome-wide in the PNUTS mutant. However, we also detected accumulation of transcripts at dSSRs, illustrated by the genome-wide map of transcript levels (S6 Fig) and the metaplot of dSSRs (Fig 4C and S7 Fig). At these divergent TSSs, the region upstream of the start site produced more transcripts in the PNUTS mutant when forward and reverse reads were analyzed separately, indicating that transcription initiates upstream of its start site upon the loss of PNUTS. In some cases, this leads to expression of genes present in the dSSR that are silent in WT cells. This explains the mapping of genes upregulated in the PNUTS mutant to 5’-end PTUs (Fig 4A). A specific example, shown in Fig 5A, includes a dSSR on chromosome 10 where a gene (Tb927.10.8340) located between the two divergent TSSs (mapped by tri-phosphate RNA sequencing of WT cells) is affected by the loss of PNUTS. Specific upregulation of this gene 4- to 13-fold following the loss of PNUTS, JBP3 and Wdr82 versus genes in the adjacent PTU is confirmed by RT-qPCR (Fig 5B). Another example confirmed by RT-qPCR is shown in S8 Fig where the gene (Tb927.10.6430) located upstream of the TSS in WT cells is specifically upregulated 4- to 7.5-fold by the loss of PNUTS, JBP3 and Wdr82.

**Fig 5.**
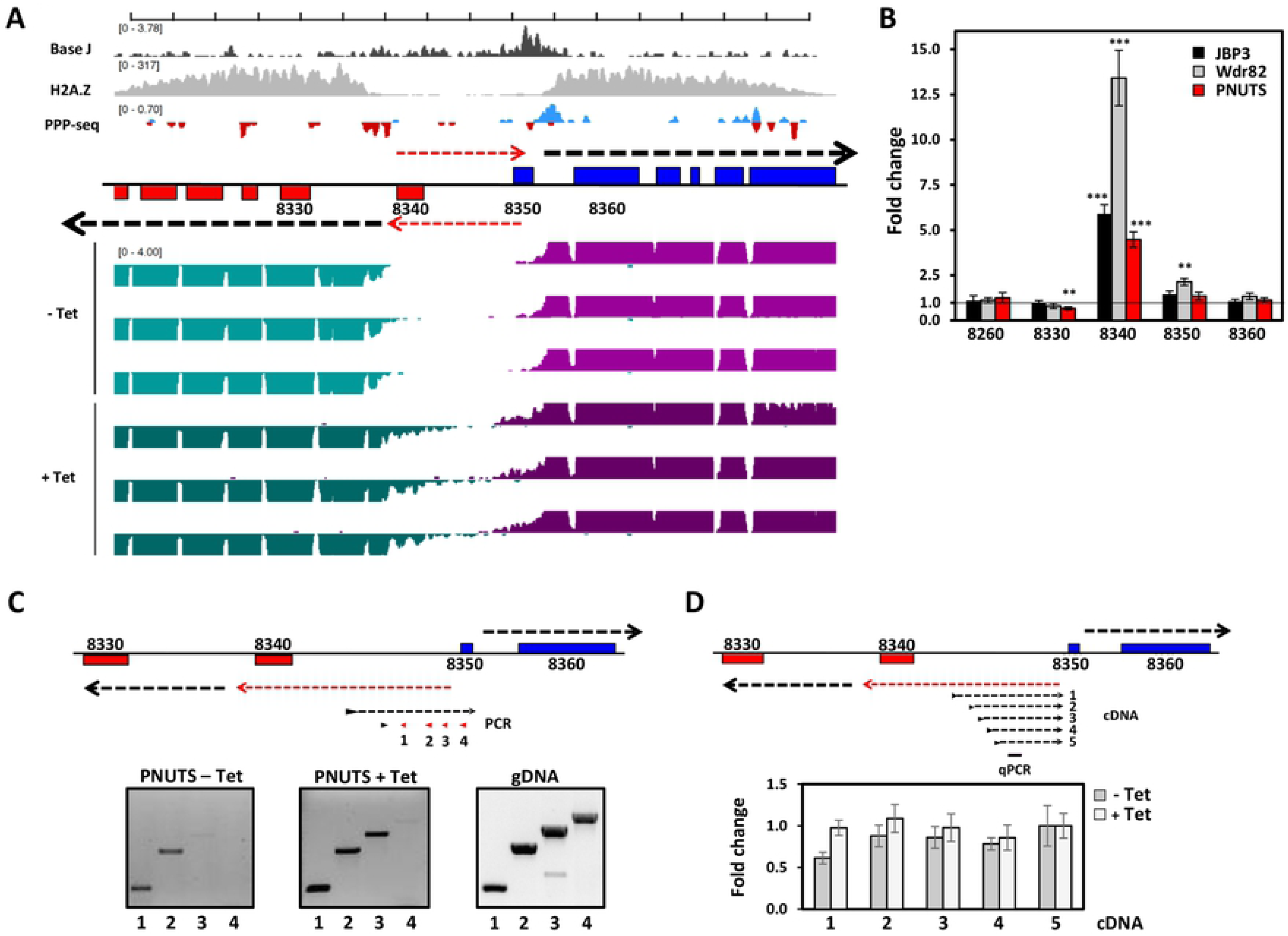
TbPNUTS affects early termination of antisense transcription at TSSs. (A) Representative region of chromosome 10 illustrating bi-directional transcription at TSSs upon TbPNUTS ablation. TSSs are denoted by PPP-seq and H2A.Z ChIP-seq enrichment in wild-type *T. brucei*. PPP-seq track colors: Red, reverse strand coverage; blue, forward strand coverage. RNA-seq track colors: Green, reverse strand coverage; Purple, forward strand coverage. Black arrows indicate direction of sense transcription. Red arrows indicate stimulated antisense transcription in the PNUTS mutant that leads to de-repression of the annotated 8340 gene on the bottom strand and, to a lesser degree, the 8350 gene on the top strand. (B) Confirmation of mRNA-seq transcript changes in the PNUTS, Wdr82 and JBP3 RNAi by RT-qPCR. RT-qPCR analysis was performed for the indicated genes as described in Material and Methods. Transcripts were normalized against Asf1 mRNA, and are plotted as the average and standard deviation of three replicates. P values were calculated using Student’s t test. **, p value ≤ 0.01; ***, p value ≤ 0.001. (C) Mapping of the 5’ end of the antisense transcript by nested RT-PCR on strand-specific cDNA. PCR utilized constant 3’ primer (indicated by black arrow-head) with varying 5’ primers indicated in red (primers 1-4). Transcript levels were normalized against strand-specific Asf1 mRNA. (D) Strand-specific RT-qPCR analysis of antisense nascent transcript. cDNA was generated using various strand specific 3’ primers and RNA from – and + Tet. qPCR was then done using internal primers to amplify the region indicated by black bar. Transcript levels were normalized against strand-specific Asf1 mRNA. Error bars indicate standard deviation from at least three experiments.

Many promoters for RNA Pol II are bidirectional in organisms from yeast to human (61–65). Unidirectional transcription resulting in productive mRNAs is typically ensured since antisense transcription is susceptible to early termination linked to rapid degradation. Mapping TSSs using tri-phosphate RNA-seq has previously suggested bi-directional activity, with strong strand bias, of Pol II initiation sites in *T. brucei* (66, 67). The RNA-seq data presented here is consistent with bi-directional activity of TSS, early termination and stimulation of divergent antisense transcription in the PNUTS knockdown leading to the activation of genes within the dSSR (Fig 4C and 5A and S8). As shown in Fig 5A and B, the silent 8340 gene within a dSSR on chromosome 10 is transcribed upon the loss of TbPNUTS. To see whether there is a correlation between the initiation of the antisense transcript with sense mRNA coding strand TSS, we analyzed the 5’-end of the nascent antisense transcript at this dSSR using RT-PCR (Fig 5C). The significant drop of inability of a 5’ primer to amplify the cDNA corresponding to the nascent ‘antisense’ transcript indicates the 5’ end of the PNUTS regulated transcript is adjacent to the TSS for the sense transcript on the opposing strand. Using various 3’ primers for generating cDNA, antisense transcription is attenuated in uninduced cells as shown in the decline in cDNA with increasing length of the transcript (Fig 5D). In the absence of PNUTS the antisense transcription fails to terminate, as seen in the maintained levels of cDNA for all primers tested in + Tet. The increasing effects of the loss of PNUTS on level of transcript with increasing length of the transcript supports the idea of PNUTS regulating early termination/elongation of the antisense transcript. Taken together the results suggest that the PJW/PP1 complex (PNUTS, JBP3 and Wdr82) regulates premature termination of antisense transcription from bi-directional TSS and silencing of gene expression in *T. brucei*.

Another possibility is that increased transcription within dSSRs is due to transcription initiating upstream of the divergent TSSs in the absence of PNUTS. To address this possibility, we examined transcription at Head-to-Head (HT) boundaries of PTUs. HT sites are defined where transcription of one PTU terminates and transcription of another PTU on the same DNA strand initiates. In contrast to TSSs at dSSRs, HT sites contain a termination site for the upstream PTU, indicated by H3.V and H4.V, and a TSS marked by individual peaks of histone variants, such as H2A.Z, and histone modifications (22, 66, 67). Similar to the metaplot analyses of dSSRs and cSSRs, we detected accumulation of transcripts at HT sites upon the ablation of PNUTS (Fig 4D). At these non-divergent TSSs, the region upstream of the start site produced more transcripts in the PNUTS mutant indicating readthrough transcription, as expected, or possible initiating events further upstream. More interestingly, antisense transcripts are also increased at HT sites. The lack of a divergent TSS at these sites, indicated by the presence of single peak of H2A.Z, supports the conclusion that the antisense transcripts are the result of regulated bi-directional transcription activity.

### Replication is not affected by pervasive transcription

Noncoding transcription, via defects in transcription termination, influences eukaryotic replication initiation. Transcription through origins located at 5’- and 3’-ends of Pol II transcription units leads to replication defects via dissociation of the prereplication complex (pre-ORCs) or sliding of MCM helicases (68–72). Of the 40 early firing origins that have been mapped in *T. brucei*, 36 are upstream of TSS (73). Analysis of *L. major* has indicated that replication initiation sites occur at the genomic locations where Pol II stalls or terminates, including sites precisely downstream of base J (74). Therefore, increased transcription at PTU flanking regions in the absence of TbPNUTS may cause DNA replication defects. To see whether pervasive transcription has any effect on *T. brucei* replication, we first analyzed whether TbPNUTS is required for proper cell-cycle progression. In *T. brucei*, DNA replication and segregation of kinetoplastid DNA (K) in the single mitochondrion precede those of nuclear DNA (N), so cells at different stages can be distinguished by their N and K configurations. 1N1K content indicates that cells are in G1, 1N2K indicates cells in S phase and 2N2K indicate post-mitotic cells. Representatives of cells with these DNA contents upon DAPI staining are shown in S9A Fig. We detect no change in cell populations following a 2 day induction of PNUTS RNAi ablation. To take a closer look at DNA replication and confirm the lack of cell cycle defects we monitored cell-cycle progression after RNAi ablation by flow cytometry, staining bulk DNA with propidium iodide and newly replicated DNA by BrdU incorporation. To do this we labeled TbPNUTS RNAi cell line (treated with tetracycline for 0, 1 and 2 days) with BrdU and BrdU labeled DNA was then detected using anti-BrdU antibodies. As shown in S9B and C Fig, uninduced cells show normal cell cycle profiles. Two days after RNAi ablation, there is no change in the cell-cycle profile or quantities of cells at each stage. Similarly, there is no detectable change in levels of BrdU incorporation in the S phase population (S9C Fig). The cell cycle profiles of the conditional RNAi ablation suggest that TbPNUTS is not required for proper cell-cycle progression and DNA replication. Furthermore, the increase in pervasive transcription in the PNUTS mutant has no measurable effect on DNA replication.

### PNUTS regulates Pol I transcriptional repression of telomeric PTUs

In addition to Pol II termination sites distributed throughout the *T. brucei* genome, H3.V and base J localize within the ~14 Pol I transcribed polycistronic units located at the telomeric ends and involved in antigenic variation (so called bloodstream form VSG expression sites, BESs) (Fig 6A)(19, 75–77). Monoallelic expression of a VSG ES leads to the expression of a single VSG on the surface of the parasite, a key aspect of the strategy bloodstream form trypanosomes use to evade the host immune system. We have previously shown that the loss of H3.V and J leads to increased expression of VSG genes from silent telomeric BESs (21). This effect is presumably due to the role of these epigenetic marks in attenuating transcription elongation Pol I within the silent VSG BESs, thereby preventing the transcription of silent VSGs. Differential gene expression analysis of RNA-seq reads mapped to the 427 genome indicates that the loss of PNUTS leads to increased VSG expression from silent BESs (S2 Table). In addition to the BES, depletion of TbPNUTS results in de-repression of VSGs from the silent telomeric metacyclic ES (MES), which are transcribed monocistronically by Pol I. A few of these VSG gene expression changes have been confirmed by RT-qPCR (S10 Fig). To further explore the global function of TbPNUTS in VSG expression control, we mapped the RNA-seq reads to the VSGnome (78). The VSGnome allows the analysis of VSG genes, such as those on minichromosomes, that were not included in the new *T. brucei* 427 genome assembly. As shown in Fig 6, this analysis confirms the de-repression of Pol I transcribed VSG at BES and MES upon TbPNUTS depletion. On the other hand minichromosomal (MC) VSGs lacking a promoter (79) were not significantly affected; only 2 of 41 were upregulated (P_adj_<0.1). These data indicate that transcription of (silent) telomeric VSGs in the PNUTS mutant is strongly dependent on the presence of a Pol I promoter. Interestingly, comparing the expression level of MES VSGs that are adjacent to the promoter with BES VSGs over 10 kb downstream (Fig 6A and S10 Fig) suggested that the level of derepression is a function of distance from the Pol I promoter. As previously mentioned, the majority of VSG genes upregulated in the PNUTS mutant are chromosomal internal VSGs (S2 Table and S5 Fig). In the VSGnome, these (unknown) VSGs (Fig 6B) were thought to be primarily located at subtelomeric arrays, but their exact positioning in the genome was not known (78). These VSGs have now been mapped to the silent subtelomeric arrays assembled in the new 427 genome (77) and, as shown in S5 Fig and Fig 6B, a significant fraction are upregulated upon the loss of PNUTS. Overall, these data indicate that TbPNUTS (PJW/PP1 complex) regulates transcriptional silencing of telomeric Pol I transcribed telomeric VSG PTUs and Pol II transcription of VSGs within genome internal PTUs in *T. brucei*.

**Fig 6.**
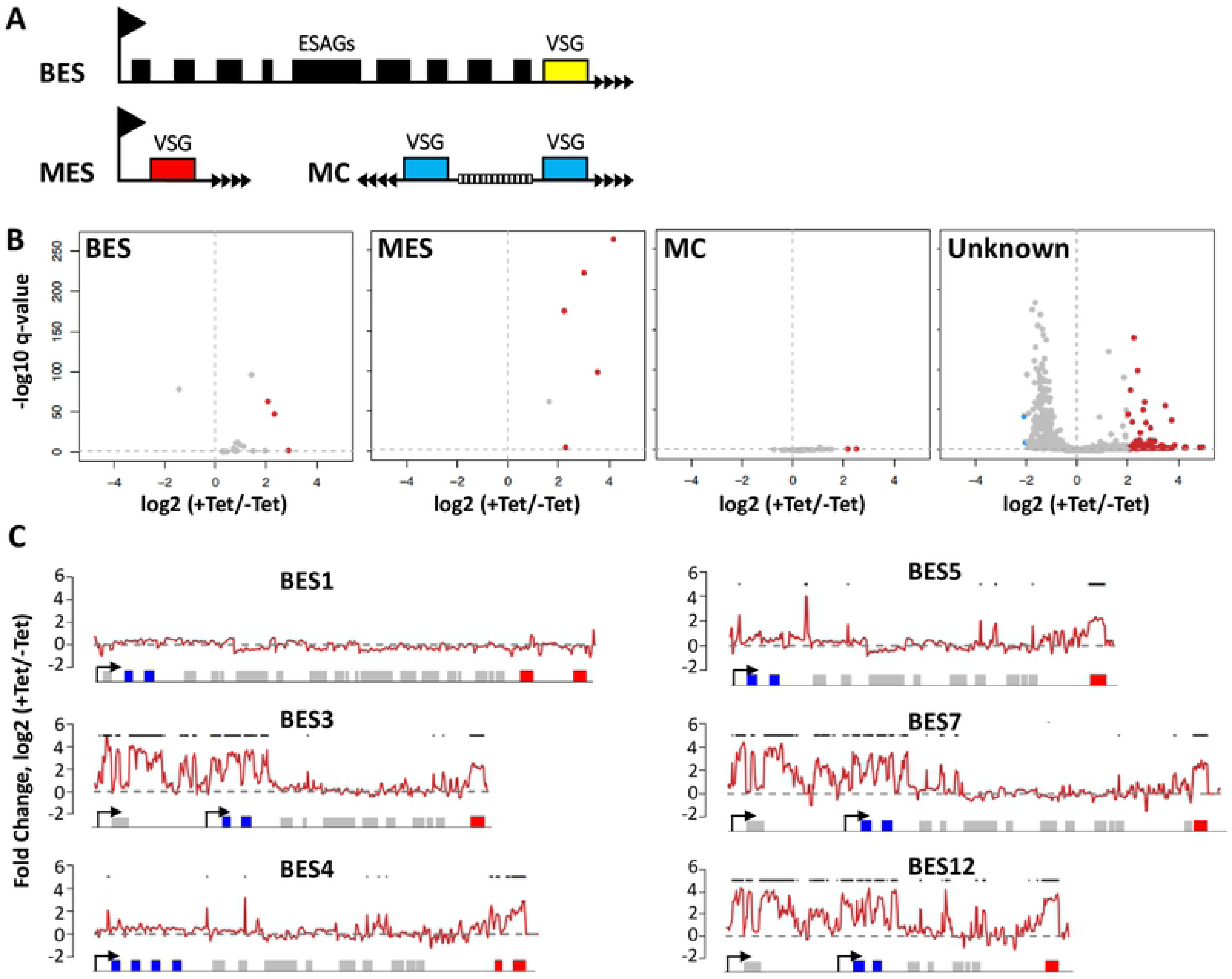
TbPNUTS is required for silencing of VSG genes. (A) Schematic diagrams of telomeric VSG genome locations; BES, Bloodstream-form Expression sites; MES, metacyclic expression sites; and MC, minichromosomal sites. ESAGs, expression site associated genes. (B) Reads from the RNA-seq experiment were aligned to the VSGnome database and raw reads mapping to each VSG was analyzed with DEseq as described in Materials and Methods. Fold changes comparing transcript levels between day 0 and day 2 following induction of PNUTS RNAi were calculated and plotted for BES, MES or MC VSGs. The rest of the VSGs (‘Unknown’) excluding BES, MC and MES were graphed separately. Red dots represent genes with greater than 3-fold change that are also significant with a Benjamini-Hochberg FDR test correction. (C) PNUTS regulates transcription of VSG BES. RNA-seq reads were aligned to the *T. brucei* 427 BES sequences (14 BESs). Fold changes comparing – and + Tet were plotted over each BES. Six of the BES are shown here. See S11 Fig for all BES. Diagram indicates annotated genes, boxes, within the BES PTU. ESAG 6 and 7 are indicated in blue. The last gene on the right is the VSG gene (red). Some BESs have pseudo-VSG genes upstream. Promoters are indicated by arrows. Some BESs have two promoters. Asterix denote bins with greater than 3-fold change of expression following PNUTS ablation.

The derepression of ESAG 6 and 7 genes adjacent to the BES promoter along with the VSG ~40 kb downstream (S2 Table), suggests that PJW/PP1 may function throughout the telomeric PTU. To examine this more closely, we analyzed the RNA-seq reads mapping to the 14 telomeric BES sequences (80). RNA-seq reads mapping to BES were counted in 200bp windows with a 100bp steps. Read counts were converted into reads per million (RPM) and compared between +/− Tet to estimate log2 fold change and plotted in Fig 7C and S11 Fig. The location of BES promoters is indicated by an arrow. The transcription of the active BES1 was not affected by the loss of TbPNUTS. However, when the remaining 13 silent BESs are analyzed we see derepression of ESAGs as well as the terminal VSG. In some cases, it seems that derepression extends 10-20 kb from the promoter to express ESAG 6 and 7, with no significant effect on the remaining genes (ESAGs) within the silent BES, and derepression of the VSG at the 3’ end. In other cases there is selective upregulation of the VSG gene, including the pseudo VSGs that are present upstream of some telomeric VSGs. The apparent selective VSG upregulation may be due to the combined effect of low level transcription of the entire BES and enhanced stability of VSG mRNAs. For example, the increased VSG mRNA half-life (4.5 hrs) compared with ESAG 6 and 7 (1.8-2.8 hrs) (81, 82). Transcripts of genes located approximately 5 kb upstream of the VSG have also been shown to be selectively rapidly degraded, presumably by nonsense-mediated decay (83). We also noticed that increased levels of RNA close to the promoter are significantly higher when there is an additional BES promoter upstream for ESAG10. In fact, the most significant gene expression changes are in BESs that have dual promoters (S2 Table). These data would suggest derepression of telomere-proximal VSG genes after PNUTS depletion is due to transcriptional activation of silent Pol I promoters. However, these results are also consistent with increased Pol I elongation along the BES. Repression of silent ESs is mediated at least in part by the inhibition of Pol I elongation within the BES preventing the production of VSG mRNA from the silent BESs (84–86). Similar to its inhibition of Pol II transcription elongation at termination sites at the 5’ and 3’ end of PTUs genome-wide, PNUTS may also function at telomeric regions to attenuate Pol I transcription elongation within the silent ESs. The data here suggest PJW/PP1 controls VSG silencing at BESs by regulating Pol I elongation (termination) and/or regulating access of the polymerase to silent promoter regions.

**Fig 7.**
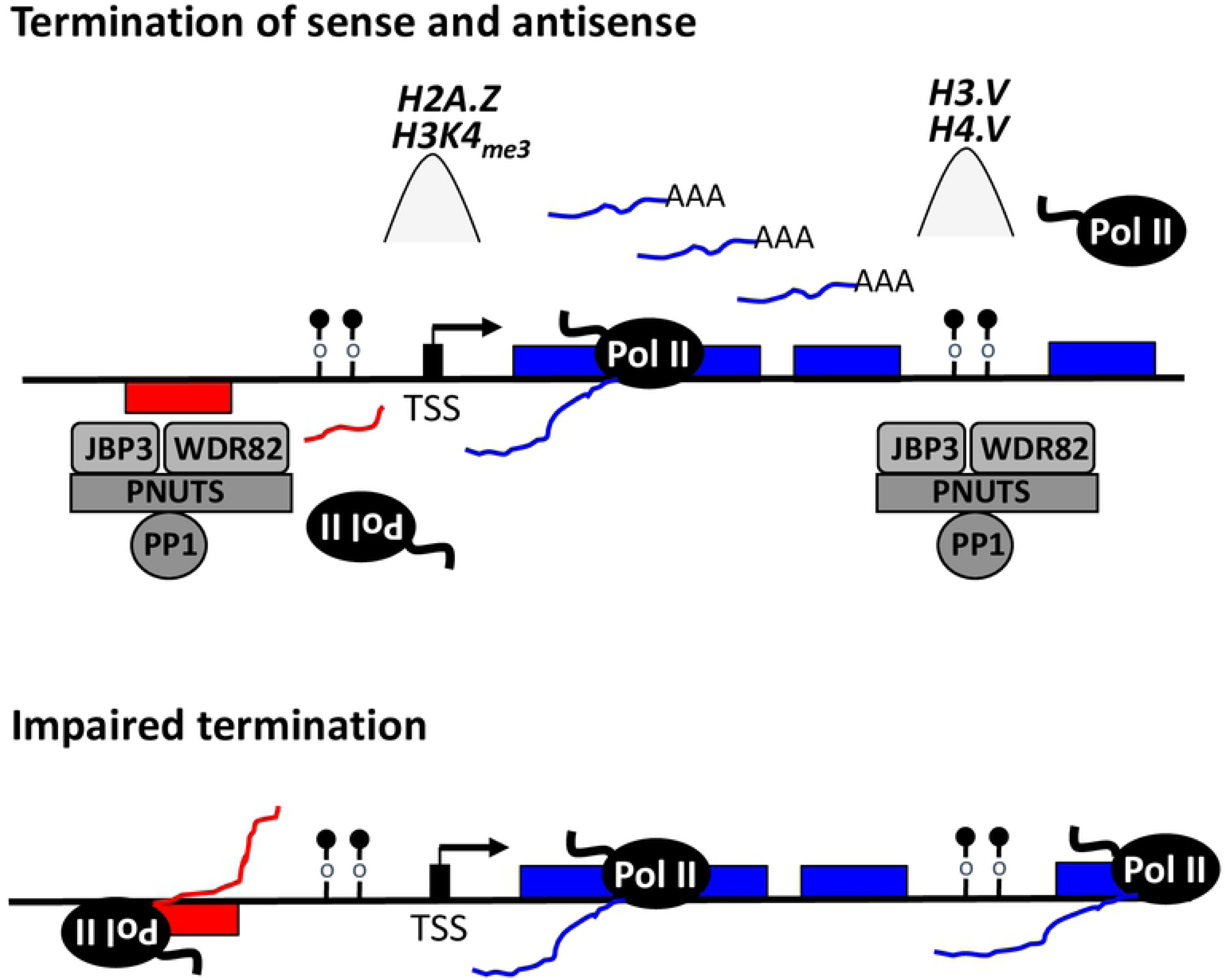
Regulation of termination by the PJW/PP1 complex in kinetoplastids. Schematic diagram of polycistronic RNA Pol II transcription. Transcription start sites (TSS), and direction of transcription, for the PTU on the top strand (blue genes) and bottom strand (red genes) are indicated by the black arrow. Peaks of histone variant (H2A.Z) and methylated histone H3 (H3K4me3) localized at TSS are indicated. According to the model, bi-directional transcription initiates at each TSS, but only the Pol II transcribing the ‘sense strand’ fully elongates and generates productive poly (A) mRNAs. Pol II terminates the 3’-end of the PTU marked with base J (ball and stick) and H3.V (not shown). The PJW/PP1 complex is recruited to the termination site via JBP3 recognition of J-DNA and termination regulated by PP1 dephosphorylation of the CTD of Pol II. The PJW/PP1 complex is also recruited to the dSSR upstream of the TSS, leading to premature termination of antisense transcription. These short transcripts may be additionally targeted for degradation. Impaired termination, following mutation of the PJW/PP1 complex, leads to readthrough transcription at the 3’- and 5’-end of PTUs. Genes located downstream of these termination sites, at the 3’- and 5’-end (upstream of the TSS) of a PTU, can generate stabilized polyA mRNAs and be expressed.

## Discussion

Current dogma in the field is that most, if not all, Pol II transcribed gene regulation is at the posttranscriptional level in kinetoplastids (87). This is primarily based on the polycistronic arrangement of genes and identification of posttransciptional regulons under the control of RNA binding protein regulated mRNA stability. Our previous studies on base J/H3.V function revealed regulated transcription termination at the 3’-end of PTUs as a novel mechanism of gene expression control in kinetoplastids. The studies described here solidify this concept and open up another possible regulatory gene expression mechanism via early termination at the 5’-end of PTUs. These studies also provide the first direct mechanistic link between base J and transcription termination in kinetoplastids by the identification of a multi-subunit protein complex involved in termination that binds base J.

Here we identify a new base J binding protein, JBP3, and show that it is part of a PP1-PNUTS-Wdr82 containing protein module. We named this module the PJW/PP1 module based upon the mammalian PTW/PP1 module involved in transcription termination (9). Mutation of its components, JBP3, Wdr82 and PNUTS, gives similar phenotypes in *T. brucei*, validating PJW is indeed a functional module. Our data strongly suggests that the kinetoplastid PJW/PP1 complex we identified here is reminiscent of human PTW/PP1 where PNUTS is the kinetoplastid functional homologue of human PNUTS and JBP3 the homologue of Tox4. However, while we demonstrate LtPP1 is a component of the Leishmania complex, the purified module in *T. brucei* lacks a PP1 homologue. PP1 is the only catalytic component of the human PTW/PP1 complex and dephosporylation by PP1 is directly involved in regulating Pol II termination (2, 3, 53, 54). Therefore, the apparent lack of PP1 association with the PNUTS, Wdr82 and JBP3 complex involved in transcription termination in *T. brucei* is surprising. MS analysis of the TbPNUTS complex is the result of a single purification from 12 L of PC *T. brucei* cells. The finding that one of three replicates of LtPNUTS purification resulted in all components of the complex except PP1 suggests PP1 may be the least stable component of the complex in *T. brucei*. Once we increase the yield of TbPNUTS purification, and reduce the volume of cells needed, we can perform multiple pulldown/MS analyses to address this possibility. We propose that PP1 is a functional component of the PJW/PP1 complex in both Leishmania and *T. brucei*, but association with TbPNUTS is less stable and/or regulated via phosphorylation. However, further work needs to be done to characterize the role of TbPNUTS phosphorylation and determine whether PP1 is a component of the PTW complex and involved in termination in *T. brucei*.

We and others have previously shown that base J and H3.V co-localize at Pol II termination sites (20, 22) and are involved in transcription termination in *T. brucei* and *L. major*, where loss of base J and H3.V leads to readthrough transcription at the 3’-end of PTUs (21, 23–26). For ‘premature’ termination sites within PTUs, read-through transcription resulting from the loss of base J and/or H3.V leads to transcription of silent genes (primarily VSG genes) downstream of the termination site (21, 24, 25). We now show that depletion of components of the PJW/PP1 complex in *T. brucei* leads to similar defects in Pol II termination at the 3’-end of PTUs, including the de-repression of downstream genes. We also uncover additional defects at the 5’-end of PTUs suggesting regulated early termination of antisense transcription from bi-directional transcription start sites. Here, we propose a model where divergent transcription at the 5’-end and readthrough transcription at the 3’-end of gene arrays is affected by the PJW/PP1 complex in trypanosomatids (model in Fig 7). According to this model, the PJW/PP1 complex is recruited to termination sites, at least partially, due to base J-JBP3 interactions. H3.V localized at 3’-end termination sites may play an additional role in complex localization since WDR5, a homolog of Wdr82, binds to the N-terminal tail of histone H3 (88). Wdr82 is required for recruitment of the APT termination complex containing PNUTS-PP1 to snoRNA termination sites in yeast (89). Wdr82 may also play a role in 5’-end termination site recognition since it binds to RNA Pol II CTD phosphorylated at Ser5 in yeast and humans (10, 90).

Similar to the mammalian complex, we propose PNUTS is a scaffolding protein for the entire PJW/PP1 complex and regulates PP1 function via the PP1 binding RVxF motif. Only three substrates have been identified for PNUTS/PP1: the Pol II elongation factor Spt5, the CTD of Pol II and MYC (2, 53, 91, 92). MYC dephosphorylation by PP1 regulates chromatin binding and stability. PP1 dephosphorylation of Spt5 and Pol II has been directly implicated in regulating Pol II termination in other eukaryotes. Therefore, we propose that regulated phosphorylation of Spt5 and Pol II by the PJW/PP1 complex is critical for transcription termination in trypanosomatids. TbSpt5 has recently been shown to be associated with Pol II (93) and is phosphorylated at a single Ser residue (94). The CTD of Pol II in trypanosomatids is unique in that it does not contain the heptad or other repetitive motifs that are conserved from yeast to humans (95). However, the CTD was shown to be essential for Pol II transcription in *T. brucei* (96, 97) and 17 phosphorylated sites have been identified within the CTD (94). Studies also suggest that CTD phosphorylation is required for Pol II association with trypanosome chromatin (98). It is necessary to determine the role of the PJW/PP1 complex in the phosphorylation status of Spt5 and Pol II in trypanosomatids and whether phosphorylation of a non-canonical Pol II CTD is involved in transcriptional regulation in divergent eukaryotes thought to lack significant regulation.

An unexpected finding of this work was that, in the absence of PNUTS, Wdr82 or JBP3, genes located upstream of TSS are also now expressed, resulting from transcription between diverging PTUs. Mammalian and yeast promoters frequently give rise to transcription in both the sense and divergent antisense directions (61–65), giving rise to a (productive) sense transcript and a corresponding upstream noncoding RNA (ncRNA) (99–102). Unidirectional transcription is typically ensured since the ncRNAs are susceptible to early termination linked to rapid degradation (100, 103–105). Early termination of divergent transcription at 5’ ends of mammalian genes occurs by similar mechanisms as termination at 3’ ends. In addition to regulating termination at the 3’ end of genes, Wdr82 and PNUTS have also been shown to be involved in enforcing early transcription termination at bi-directional promoters (17). The role of PNUTS is thought to reflect the essential nature of PNUTS/PP1 since differential phosphorylation of the CTD of Pol II has been proposed to regulate the directionality of transcription at bi-directional promoters. Specifically, Tyr1 and Ser1 hyperphosphorylation of Pol II have been shown to be associated with antisense divergent transcription at mammalian promoters (106–108). Spt5 regulation of Pol II elongation is involved in control of divergent antisense transcription as well as readthrough transcription at 3’ end of genes in yeast (2, 109), representing an additional target of PNUTS/PP1 regulation of transcription at both ends of genes. Similar to mammalian promoters where transcription is divergent and initiation is over a broad genomic region, previous studies have suggested that Pol II transcription initiation sites are intrinsically bi-directional in *T. brucei* (66, 67). We found that loss of PNUTS, Wdr82 and JBP3 can have a major effect on divergent ‘antisense’ transcription, presumably reflecting the role of PJW/PP1 in regulating termination of antisense transcription from bi-directional promoters in *T. brucei* (Fig 7). Interestingly, decreased levels of base J in Leishmania led to detection of antisense RNAs corresponding to similar regions within divergent PTUs (23). A possible explanation for antisense transcription at divergent TSSs is increased chromatin accessibility in the SSR, resulting in alternative TSS usage further upstream. However, the detection significant antisense transcription at non-divergent TSSs (HT sites) upon the loss of PNUTS strongly supports the involvement of the PJW/PP1 in regulating early termination of antisense transcription in *T. brucei*. Additional work is required to unambiguously confirm that the increased nascent antisense RNA, and corresponding mRNAs, we detect from regions upstream of TSSs are in fact a result of bi-directional transcription activity.

Loss of J and H3.V resulted in similar readthrough defects at the 3’-end of PTUs (including telomeric BESs) and gene expression as seen here in PJW/PP1 mutants, but without cell growth effects. We concluded that readthrough transcription at 3’-end and corresponding gene expression changes in the J/H3.V mutant are not lethal to the parasite. PJW/PP1 mutants we analyze here lead to additional effects on transcription and gene de-repression at 5’-ends and decreased cell growth. The ability of the PJW/PP1 complex to bind base J is consistent with its function at both ends of PTU’s. However, it is unclear why the effects in the J mutant are limited to the 3’-end and whether specific function of the complex at the 5’-end can explain its essential nature. While J is present at both 5’ and 3’ PTU flanking regions and involved in transcription, it is apparently not the dominant mark since H3.V had more significant effects on 3’-end transcription and gene expression (21). H3.V is limited to termination sites at the 3’ends of PTUs and not localized at TSSs in *T. brucei*. Furthermore, additional factors may be involved in recruitment of PJW/PP1 complex to TSS, such as Pol II-Wdr82 interactions, as mentioned above, and modified and variant histones such as H2A.Z and H3K4Me3 (Fig 7). These points may explain why PJW/PP1 mutants lead to defects at both 5’- and 3’-ends and the J/H3.V mutants are limited to the 3’-end. We propose that the essential nature of the PJW/PP1 complex is due to regulated expression of ncRNAs and/or mRNAs at the 5’-end of PTUs or additional unknown functions of JBP3, Wdr82 and PNUTS in *T. brucei*.

The possibility that the essential nature of the complex is due to regulation of transcription-replication conflicts at TSSs was directly addressed. We expected the pervasive transcription phenotype in the PJW/PP1 mutant to negatively impact DNA replication and explain the reduced cell growth. Replication origins tend to localize after TTS in yeast and upstream of promoters in humans, presumably to minimize transcription-replication conflicts (71, 110, 111). The induction of transcription through origins, via defects in transcription termination at TTSs as well as at TSSs, leads to replication defects via dissociation or sliding of the pre-ORCs and MCMs (68–72) and changes in chromatin structure (112). Furthermore, loss of origin function activates readthrough transcription in mammals and yeast. Clearly there is a relationship between transcription and DNA replication in eukaryotes. This relationship appears to exist in *T. brucei* as well, since origins flank the PTUs at TSS and TTS, and TbORC1 and TbMCM-BP deletions led to similar defects in transcription at PTU flanks as we illustrate here (73, 113). In the recent analysis of the TbMCM-BP mutant (113), in addition to its role in termination at 3’ ends, the antisense RNA at H-T sites suggested a role of TbMCM-BP in determining the direction of transcription, what we refer to as bi-directional activity, while at dSSRs they concluded it was due to alternative initiation events. Regardless of the mechanism involved, increased transcription upstream of TSSs in the PJW/PP1 mutants would presumably result in defects in replication. However, the TbPNUTS mutant does not indicate any alteration in cell cycle or DNA replication. Alternatively, the level of transcription induction at origins is too low at day two of the RNAi for any significant effects on replication. Further work is needed to explore the effects of increased transcription on ORC function in trypanosomes.

Functional interaction between replication and transcription machineries was further suggested by derepression of Pol I transcribed silent BESs and MESs in the TbORC1 and TbMCM-BP mutants (73, 114). How these operate mechanistically at ESs, regulating Pol I elongation via chromatin changes along silent BESs or BES promoter activity, remains unclear. In fact, whether VSG monoallelic expression control of BESs takes place at the initiation or elongation level is still debated. Low levels of transcripts from silent ESs upon the knockdown of factors such as ORC1, MCM-BP and PNUTS cannot resolve this issue, since the data are compatible with both models. While derepression of the BES in the PNUTS mutant would suggest the PJW/PP1 complex has a direct effect on the activity of silent BES promoters, the proposed role of the complex in regulating Pol II elongation/termination at 5’ and 3’ end of PTUs genome-wide would suggest a similar role in regulating Pol I elongation. If so, presumably Pol I is regulated in a different manner, since there is no evidence for phosphorylation of Pol I. However, the Spt5 substrate for PNUTS-PP1 has been shown to bind and regulate Pol I transcription in mammalian cells (115, 116). Further studies are necessary to understand how PJW/PP1 complex regulation of Pol II transcription at 5’ and 3’ ends of PTUs genome-wide is related to silencing of the specialized telomeric Pol I PTUs. For example, derepression of BES promoters may be a functional consequence of non-coding RNA transcription generated by pervasive Pol II transcription in the PNUTS mutant. We have previously shown that readthrough transcription at 3’ ends of PTUs lead to significant levels of siRNAs in *T. brucei* (21). Regardless of the mechanism involved, the complex is required not only for repression of telomeric and subtelomeric VSGs but also VSGs scattered within the chromosome at 3’ ends of PTUs. Depletion of PNUTS also increased the levels of procyclin and PAG RNAs, which are transcribed from a Pol I promoter that is repressed in BSF *T. brucei*. This transcription unit is located at the end of Pol II transcribed PTU and increased transcription in PNUTS mutant may be due to readthrough transcription as in the J/H3.V mutant (21). Thus, the PJW/PP1 complex is required for repression of life-cycle specific genes transcribed by Pol I in the mammalian infectious form of *T. brucei*. We have therefore uncovered a functional link between transcriptional termination and Pol II- and Pol I-mediated gene silencing in *T. brucei*.

## Materials and Methods

### Parasite cell culture

Bloodstream form *T. b. brucei* Lister 427 (MiTat 1.2) or “single marker cells (SMC)”, expressing T7 RNA polymerase and the Tet repressor (117) were used in these studies and cultured in HMI-9 medium. Transfections were performed using the Amaxa electroporation system (Human T Cell Nucleofactor Kit, program X-001). Where appropriate, the following drug concentrations were used: 2 μg/ml G418, 5 μg/ml Hygromycin, 2.5 μg/ml Phleomycin, 2 μg/ml Tetracycline. Procyclic form *T. b. brucei* TREU667 and promastigote form *L. tarantolae* were cultured in SDM79 medium. Transfections were performed using the BioRad GenePulser II (2 pulses at 1.4 kV / 25 μF) in 0.4 cm cuvettes with 1 × 10^8^ cells (*L. tarantolae*) or 3 × 10^7^ cells (PCF *T. b. brucei*) in 0.5 ml cytomix (2 mM EGTA, 120 mM KCl, 0.15 mM CaCl_2_, 10 mM KP_i_ pH = 7.6, 25 mM HEPES pH = 7.6, 5 mM MgCl_2_, 0.5 % glucose, 100 μg/ml BSA, 1 mM Hypoxanthine). Where appropriate, the following drug concentrations were used: 25 μg/ml G418 (PCF *T. b. brucei*) and 50 μg/ml G418, 10 μg/ml Puromycin (*L. tarantolae*).

### RNAi analysis

For conditional PNUTS, JBP3 and Wdr82 silencing experiments in *T. brucei* a part of the ORF was integrated into the BamH I site of the p2T7-177 vector (118). Sce-I linearized p2T7-177 constructs were transfected into BF SMC for targeted integration into the 177bp repeat locus. RNAi was induced with 2 μg/ml Tetracycline and cell growth was monitored daily in triplicate. Primers sequences used are available upon request.

### Epitope tagging of proteins

For generation of the dual (Protein A and Streptavidin Binding Protein) tagged constructs in *L. tarantolae*, the coding regions of LtJGT, LtJBP3, and LtWdr82 lacking stop codons were amplified and cloned into the BamHI and XbaI sites of pSNSAP1 (42). For PTP tagging in *T. brucei*, the 3’ end of *T. brucei* genes were cloned in the Apa I and Not I sites of the pC-PTP-Neo vector (52), resulting in dual (Protein A and Protein C) tagged proteins. Linearization of the constructs was performed using a unique restriction site within the 3’ end of the cloned gene. All final constructs were sequenced prior to electroporation. Tagging the 3’-end of the TbPNUTS, TbJBP3 and TbWdr82 with 3x HA tag was performed using a PCR based approach with the pMOTag4H construct (119). Primers sequences used are available upon request.

### Tandem affinity purification (TAP) and co-immunoprecipitation

Tandem affinity purification was performed from whole cell extracts. 2 × 10^11^ cells (*L. tarantolae* and PCF *T. brucei*) were harvested, 1 time washed in 1 × PBS (137 mM NaCl, 2.7 mM KCl, 8 mM Na_2_HPO_4_, 2 mM KH_2_PO_4_ pH = 7.4), 2 times washed in buffer I (20 mM Tris pH = 7.7, 100 mM NaCl, 3 mM MgCl_2_, 1 mM EDTA) and 1 time washed in buffer II (150 mM Sucrose, 150 mM KCl, 3 mM MgCl_2_, 20 mM K-glutamate, 20 mM HEPES pH = 7.7, 1 mM DTT, 0.2 % Tween-20). Cell pellets were then adjusted to 20 ml (*L. tarentolae*) or 50 ml (PCF *T. brucei*) with buffer II with protease inhibitors (8 μg/ml Aprotinin, 10 μg/ml Leupeptin, 1 μg/ml Pepstatin, 1 mM PMSF and 2 tablets cOmplete Mini, EDTA free; Roche) and 2 times flash frozen in liquid Nitrogen. Lysates were then sonicated (Sonics, Vibra-Cell) for 5 times (15” on / 45” off, 50% amplitude, large tip) on ice. Extracts were cleared by centrifugation for 10 min at 21,000 × g at 4°C and incubated while rotating with 200 μl IgG Sepharose beads (GE Healthcare) for 4 hrs at 4°C. The beads were then washed with 35 ml PA-150 buffer (20 mM Tris pH = 7.7, 150 mM KCl, 3 mM MgCl_2_, 0.5 mM DTT, 0.1 % Tween-20) and 15 ml PA-150 buffer with 0.5 mM EDTA. The beads were then resuspended in 2 ml PA-150 buffer with 0.5 mM EDTA and 250 U TEV Protease (Invitrogen) while rotating for 16 hrs at 4°C. The supernatant was collected and for *L. tarantolae*, samples were incubated with 100 μl magnetic Streptavidin C1 Dynabeads (Invitrogen) for 4 hrs while rotating at 4°C. Beads were then washed with 50 ml PA-150 buffer, transferred into 1 ml elution buffer (100 mM Tris pH = 8.0, 150 mM NaCl, 1 mM EDTA, 2.5 mM *d*-desthiobiotin), incubated for 2 hrs while rotating at room temperature. Eluted protein was then TCA precipitated and subjected to MS/MS analysis. To the *T. brucei* supernatant samples, CaCl_2_ was added to a final concentration of 1.25 mM and incubated with 200 μl Anti-Protein C Affinity Matrix (Roche) for 4 hrs while rotating at 4°C. Beads were then washed with 50 ml PA-150 buffer with 1.25 mM CaCl2. Bound protein was eluted in 5 ml elution buffer (5 mM Tris pH = 8.0, 5 mM EDTA, 10 mM EGTA), TCA precipitated and subjected to MS/MS analysis.

Pull-down experiments using PTP-tagged PNUTS, JBP3 and Wdr82 were performed using the first purification step of the TAP protocol on whole cell extracts from 2 × 10^8^ cells in 200 μl buffer II. After the IgG Sepharose incubation, beads where washed in PA-150 buffer and boiled in Laemmli buffer. For Western analysis, 5 × 10^6^ cell equivalents were analyzed on 10% PAA / SDS gels and sequentially probed with anti-HA antibodies (Sigma, 3F10, 1:3000), anti-Protein A antibodies (Sigma, P3775, 1:5000) and anti-La antibodies (a gift from C. Tschudi, 1:500).

### MS-MS analysis

Eluted proteins were digested in-solution as previously described (120). Briefly, eluted proteins were reduced with dithiothreitol (DTT) at 56°C for one hour followed by carboxyamidomethylation with 55mM iodoacetamide in the dark for 45 minutes, and finally digested with sequencing grade trypsin overnight. Protease activity was halted by the addition of trifluoroacetic acid and resulting tryptic peptides were desalted using C18 spin columns (The Nest Group, SUM SS18V). Following clean up, peptides were dried down and reconstituted with 5% acetonitrile in 0.1% formic acid and spun through a 0.2um bio-inert membrane tube (PALL, ODM02C35) prior to autosampler tube transfer. Peptides were analyzed on an Orbitrap Fusion Lumos Tribrid mass spectrometer (Thermo Fisher Scientific) interfaced with an UltiMate 3000 RSLCnano HPLC system (Thermo Scientific). Peptides were resolved on an Acclaim™ PepMap™ RSLC C18 analytical column (75um ID × 15cm; 2µm particle size) at a flow rate of 200nL/min using a gradient of increasing buffer B (80% acetonitrile in 0.1% formic acid) over 180 minutes. Data dependent acquisition was carried out using the Orbitrap mass analyzer collecting full scans every three seconds (300-2000 m/z range at 60,000 resolution). The most intense ion that met mono-isotopic precursor selection requirements were selected, isolated, and fragmented using 38% collision-induced dissociation (CID). Every precursor selected for ms/ms analysis was added to the dynamic exclusion list and precursors selected twice within 10 seconds were excluded for the following 20 seconds. Fragment ions were detected using the ion trap to increase the duty cycle and achieve more ms/ms scans per experiment. All raw ms/ms spectra were searched against the GeneDB *Leishmania tarentolae* and T. *brucei* 927 database using the Seaquest search algorithm in Proteome Discoverer 2.2 (Thermo Fisher Scientific). Search parameters were set to allow for two missed tryptic cleavages, 20 ppm mass tolerance for precursor ions, and 0.3 Daltons mass tolerance for fragment ions. A fixed modification of carbamidomethylation on cysteine residues and a variable modification of oxidation on methionine residues were enabled to accurately match fragment ions. A fixed value peptide spectral match (PSM) validation node was used to validate each PSM at a maximum delta Cn of 0.05. All PSMs were subjected to further validation with a maximum of 1% false discovery rate utilizing PSMs matched to a reverse (decoy) database while being assigned to protein groups.

### 3D structure prediction of JBP3

To analyze the putative J-DNA binding domain of JBP3, the representative region of LtJBP3 was threaded through the JBP1 JBD PDB structure to search for similar secondary structural folds using the comparative modeling program I-TASSER (iterative threading assembly refinement algorithm). (http://zhang.bioinformatics.ku.edu/I-TASSER) The initial 3D models generated by I-TASSER were of high quality, with a C-score of −0.76 and a TM-score of 0.62 (within the correct topology range of I-TASSER). TM-score is defined to assess the topological similarity of the two protein structures independent of size. A TM-score >0.5 corresponds approximately to two structures of the similar topology. The top predicted model was then aligned using TM-align from I-TASSER with the Lt JBD model 2xseA giving a TM-score of 0.727, RMSD of 1.46, and Cov score of 0.774.

### Recombinant protein production

The *L. tarentolae* JBP3 gene was amplified from genomic DNA by PCR with primers containing 5’ Nde I and 3’ BamH I restriction sites. PCR fragments were digested with Nde I and BamH I and subcloned into the pet16b expression vector. Plasmids were transformed into Escherichia coli (BL21 DE3), and bacteria were cultured in defined autoinduction media to allow growing cultures to high densities and protein expression without induction, as previously described (121). Briefly, LB media is supplemented with KH_2_PO_4_, Na_2_HPO_4_ and 0.05% Glucose, 0.2% Lactose and 0.6% Glycerol. Bacteria were cultured in this media in the presence of 100μg/ml ampicillin at 37°C for 2 h and then shifted to 18°C for 24 h. Cells were lysed and JBP3 purified by metal affinity as previously described for JBP1 (37, 38). The affinity-purified JBP3 was concentrated to 0.5 ml in a Centricon-100 apparatus and loaded onto a Sephadex S-200 (Amersham Biosciences 16/60) column equilibrated with buffer A (50 mM Hepes, pH 7.0, 500 mM NaCl, 1 mM DTT). The fraction containing JBP3 was concentrated to 200 ul by Centricon-100. Protein purity was analyzed by SDS-PAGE stained with Coomassie Brilliant Blue, and protein concentration was determined using BSA standards.

### J-DNA binding

Electrophoretic Mobility Shift Assays were carried out essentially as described previously for JBP1 and VSG-1J DNA substrate (37, 38), with few changes. The binding reaction mixture (20 ul) contained 35 mM Hepes-NaOH, pH 7.9, 1mM EDTA, 1 mM DTT, 50 mM KCL, 5 mM MgCl, 10 ug BSA and 15 fmol radiolabeled DNA substrate containing a single modified J base (VSG-1J) or no modified base, and indicated JBP3 protein amounts. The reactions were incubated for 15 min at room temperature and analyzed on a 4.5% nondenaturing polyacrylamide gel at 4°C. After drying, the gels were exposed to film.

### Sucrose sedimentation analysis

For the sedimentation analysis of the PJW/PP1 complex, extracts were made from the BF *T. brucei* cell lines expressing PNUTS-PTP and JBP3-HA or Wdr82-HA and loaded onto 10 ml, 5-50% sucrose gradients. Samples were centrifuged at 38000 rpm for 18 hours using a SW41 Ti rotor (Beckman). The gradient was fractionated from top to bottom in twenty aliquots of 500ul each. Proteins from each fraction were enriched by methanol: chloroform protein precipitation and resuspended in SDS loading buffer for electrophoresis.

### RT-PCR analysis

Total RNA was isolated with Tripure Isolation Reagent (Roche). cDNA was generated from 0.5 – 2 μg Turbo™ DNase (ThermoFisher) treated total RNA with Superscript™ III (ThermoFisher) according to the manufacturer’s instructions with either random hexamers, oligo dT primers or strand specific oligonucleotides. Strand specific RT reactions were performed with the strand specific oligonucleotide and an antisense Asf I oligonucleotide. Equal amounts of cDNA were used in PCR reactions with Ready Go Taq Polymerase (Promega). A minus-RT control was used to ensure no contaminating genomic DNA was amplified. Primer sequences used in the analysis are available upon request.

### Quantitative RT PCR analysis

Total RNA was isolated and Turbo™ DNase treated as described above. Quantification of Superscript™ III generated cDNA was performed using an iCycler with an iQ5 multicolor real-time PCR detection system (Bio-Rad). Triplicate cDNA’s were analyzed and normalized to Asf I cDNA. qPCR oligonucleotide primers combos were designed using Integrated DNA Technologies software. cDNA reactions were diluted 10 fold and 5 µl was analyzed. A 15 µl reaction mixture contained 4.5 pmol sense and antisense primer, 7.5 µl 2X iQ SYBR green super mix (Bio-Rad Laboratories). Standard curves were prepared for each gene using 5-fold dilutions of a known quantity (100 ng/µl) of WT gDNA. The quantities were calculated using iQ5 optical detection system software. Primers sequences used are available upon request.

### Strand-specific RNA-seq library construction

For mRNA-seq, total RNA was isolated from *T. brucei* RNAi cultures grown in presence or absence of tetracycline for two days using TriPure. Six mRNA-seq libraries were constructed for PNUTS RNA (triplicate samples for plus and minus tetracyclin) and four libraries for VAa RNA (duplicate samples) were constructed using Illumina TruSeq Stranded RNA LT Kit following the manufacturer’s instructions with limited modifications. The starting quantity of total RNA was adjusted to 1.3 µg, and all volumes were reduced to a third of the described quantity. High throughput sequencing was performed at the Georgia Genomics and Bioinformatics Core (GGBC) on a NextSeq500 (Illumina).

### RNA-seq analysis

Raw reads from mRNA-seq were first trimmed using fastp with default settings (v0.19.5; (122)). Remaining reads were locally aligned to the recently published long-read *T. brucei* Lister 427 2018 version 9.0 genome assembly (downloaded from (78)) and the Lister 427 BES sequences (80), using Bowtie2 version 2.3.4.1 (123). With non-default settings (sensitive local) and further processed with Samtools version 1.6 (124). For each sample, HTSeq (v0.9.1) was used to count reads for each reference transcript annotation, followed by normalization/variance stabilization using DESeq2 (v1.18.1). Differential gene expression was conducted using DESeq2 by comparing TbPNUTs RNAi samples with and without tetracyclin in triplicate (log2 fold change and differential expression test statistics can be found in Table S1). Due to incomplete gene annotation of the BES in the new *T. brucei* Lister 427 2018 genome assembly, gene expression changes for BES were determined by aligning reads to the Lister 427 BES sequences (S2 Table). To compare tetracyclin-treatment fold changes for specific strands genome-wide, we counted reads from each strand in 200bp bins with a 100bp step. Mapping of differentially upregulated genes in a genome-wide context was determined by highlighting genes upregulated >3-fold in red for all 11 megabase chromosomes. Fold changes between Tet-untreated and Tet-treated PNUTS RNAi were plotted for all 11 megabase chromosomes as well. Tag counts in 200bp bins (100bp steps) were used to estimate correlations among samples (correlation coefficients among replicates were >0.99).

To analyze transcription defects at 3’- and 5’-end of PTUs, reads mapping to TTS (cSSRs) and TSS (dSSRs) were counted and reads per kilobase per million mapped reads (RPKM) values were generated. Similar to what we have previously done (21, 24, 25), lists of cSSRs and dSSRs were generated computationally as defined regions where coding strands switch based on the transcriptome as well as TSS mapped via triphosphate RNA sequencing. Head-to-Tail (HT) sites were defined were one PTU terminates (H3.V) and another PTU on the same strand initiates. The TSS at HT sites were further defined by a single peak of H2A.Z and the lack of an annotated gene on the antisense strand, distinguishing it from a TSS at a dSSR. Several SSRs located at subtelomeres were not included due to ambigious nature of gene organization. SSRs and the 5-kb flanking regions were analyzed with DeepTools (v3.2.1) using 100bp bins flanking SSRs and dividing each SSR into 50 equally sized bins. Violin plots were generated using the R package vioplot, with the median and interquartile range illustrated by white circles and black boxes, respectively. Genes were considered adjacent to base J if the gene, according to the *T. brucei* Lister 427 annotation, was within 1-kb either upstream or downstream base J peaks. J IP-seq data shown here are from previously published work (20, 22).

To examine VSG expression, trimmed reads were bowtie-aligned to the VSGnome (retrieved from http://tryps.rockefeller.edu) (78) and differential expression was analyzed using the DEseq2 identically to the analysis of the entire genome.

### BrdU pulse labeling and flow cytometry

BrdU pulse labeling experiments were performed as described (125) with some modifications. Briefly, 5 × 10^7^ BSF cells were harvested, 2 x washed in serum free HMI9, supplemented with 1 % BSA and 1 % glucose, resuspended at 1 × 10^7^ / ml and pre-incubated at 37°C for 10 min. BrdU was added to a final concentration of 0.5 mM and incubated for 40 min at 37°C. Cells were then washed 2 x in PBS and resuspended at 1 × 10^7^ / ml. Ice cold 100 % ethanol was slowly added while shaking to a final concentration of 25 % ethanol. 2 × 10^7^ cells were then incubated for 15 min in 1 ml 0.1 n HCl / 0.5 % Triton X-100, washed 2 times with PBS / 1% BSA / 0.5 % Tween 20, resuspended at 2 × 10^7^ / ml and incubated with 2 µg / ml anti BrdU antibodies (Sigma B8434) for 2 hrs at RT. Cells were 1 x washed and incubated with 8 µg / ml Alexa488 labelled anti Mouse IgG (ThermoFisher, A11001) for 1 hr at RT in the dark. Cells were then washed 2 times and resuspended in 0.5 ml PBS / 1 % BSA / 1 % Tween 20 with 5 µg / ml Propidium Iodide. Analysis was performed using a Cyan cytometer (DAKO).

### Cell cycle analysis

Cells were harvested, 2 x washed in PBS, allowed to settle on glass slides and air dried for 10 min at RT. Cells were then fixed in −20°C methanol for 10 min and mounted with ProLong Gold antifade reagent with DAPI (Molecular Probes, P36935). Approximately 300 cells / sample were analyzed for their DNA content.

### Data Availability

RNA-seq raw files and processed files have been deposited to the NCBI Gene Expression Omnibus (GEO) with accession number GSE135708.

## Acknowledgments

We are grateful to Jessica Lopes da Rosa-Spiegler, David Reynolds, Nicolai Siegel, and Piet Borst for critical reading of the manuscript. We would also like to thank Shulamit Michaeli (Bar-IIan University) for the pSNSAP1 vector; Arthur Günzl (UConn Health) for the pC-PTP-NEO vector; Roberto DoCampo (University of Georgia) for VA a RNAi construct; Mike Cipriano (University of Georgia) for help with homology modeling; Christina L. Ethridge (University of Georgia) for generating the RNA-seq libraries.

## Supporting Information

**S1 Fig. JBP3 is a putative J-binding protein with a conserved JBD motif.** (A) Schematic representation of the structure of JBP1 and JBP3 from *L. tarentolae* illustrating the presence of the conserved JBD and variable C-termini. (B) A multiple sequence alignment of the JBD of JBP1 homologues from *T. brucei* (Tb927.11.13640), *L. major* (LmjF.09.1480), *T. cruzi* (TcCLB.506753.120), and *L. tarentolae* (LtaP09.1510) and the conserved region of JBP3 is shown. The sequence alignment was generated using Maft and visualized with Jalview. Identical amino acids are indicated by highlighting; >80% agreement is highlighted in mid blue and >60% in light blue. Similar amino acids are indicated by hierarchical analysis of residue conservation shown below. (C) 3D structure prediction. Using the structure of the JBD of JBP1, JBP3 was run through I-TASSER and aligns with RMSD of 1.46, Cov score of 0.774, TM-score of 0.727. In the superposition, the thick backbones are the native JBP1 JBD structure and the thin backbone is the I-TASSER model of JBP3. Blue to red runs from N- to C-terminal.

**S2 Fig. PNUTS is a disordered protein.** (A) Compositional profiling of Lt and Tb PNUTS showing the fractional amino acid composition in comparison with the compositional profile of typical ordered proteins. The compositional profile of typically disordered proteins from the DisProt database is shown for comparison below. (B) Analysis of TbPNUTS using the DISOPRED3 program for protein disorder prediction and for protein-binding site annotation within disordered regions.

**S3 Fig. Co-immunoprecipitation.** (A) Endogenously PTP tagged PNUTS, JBP3, Wdr82 and CPSF73. (A and B) Co-IP experiments as described in Fig 2A. (B) JGT is not associated with the *T. brucei* complex. (C) PP1-1, PP1-7 and HSP70 do not associate with PNUTS in *T. brucei*. (D) TbPNUTS is phosphorylated *in vivo* in BSF and PCF *T. brucei*. Extract from the indicated *T. brucei* cells expressing PNUTS-PTP were incubated with and without calf intestinal phosphatase (CIP) and western blotted using anti-prot A.

**S4 Fig. Consequence of PNUTS depletion on the *T. brucei* transcriptome.** (A) Gene expression changes upon RNAi ablation of PNUTS (left) and VA a (right) are plotted. Triplicate analysis of PNUTS and duplicate of VA a. On the left, red dots represent genes with greater than 3-fold change that are also significant with a Benjamini-Hochberg FDR test correction. On the right, red dots represent genes with greater than 2-fold change after VA a ablation in both replicates. (B) Gene expression changes at cSSRs (N=193) and dSSRs (N=197) that are within 1kb of base J matched with same number of random locations within the genome for ablation of PNUTS and VA a.

**S5 Fig. Genes that are at least 3-fold upregulated upon the loss of PNUTS are located at the 3’- and 5’-end of PTUs and subtelomeric VSG clusters.** Upregulated genes (>3-fold) upon PNUTS RNAi are indicated in red in the *T. brucei* 427 genome assembly. Only one of the two homologous chromosomes is depicted for the homologous core regions. Both chromosomes are shown for the heterozygous subtelomeric regions containing silent VSGs. The telomeric VSG expression sites are not included in this assembly. Metacyclic-form expression sites are marked with an asterisk.

**S6 Fig. Ablation of TbPNUTS accumulates antisense transcripts at PTU borders.** Transcription was measured by stranded RNA-seq. Fold changes comparing transcription levels between – and + tetracyclin induction of PNUTS RNAi were calculated in 200bp windows (100bp step) and plotted over the chromosome length. The core regions of the 11 chromosomes are shown. Forward (top strand) and reverse reads (bottom strand) were analyzed separately and plotted above and below the chromosome diagram, respectively. Blue genes are transcribed in reverse PTUs and red are forward PTUs. Asterix denote bins with greater than 2-fold change of expression following ablation.

**S7 Fig. Box plots comparing the levels of transcripts from dSSR and cSSR before and after TbPNUTS and VA a ablation.** Normalized reads per million estimates were derived for dSSRs and cSSRs as the average across replicates per sample. Median values are indicated by white dots. Differences between + and – Tet were measured by a Mann-Whitney U statistical test.

**S8 Fig. TbPNUTS affects early termination of antisense transcription at TSSs.** (A) Another divergent PTU region of chromosome 10 illustrating bi-directional transcription at TSSs upon TbPNUTS ablation. TSSs are denoted by PPP-seq and H2A.Z ChIP-seq enrichment in wild-type *T. brucei*. PPP-seq track colors: Red, reverse strand coverage; blue, forward strand coverage. RNA-seq track colors: Green, reverse strand coverage; Purple, forward strand coverage. Black arrows indicate direction of sense transcription. Red arrows indicate stimulated antisense transcription. In the case of the top strand, this ‘antisense’ transcription leads to derepression of the annotated 6430 gene (B) Confirmation of mRNA-seq transcript changes by RT-qPCR. RT-qPCR analysis was performed for the indicated genes as described in Fig 5B.

**S9 Fig. Pervasive transcription does not affect DNA replication.** (A) Cell cycle analysis of wild-type BSF *T. brucei* (WT 221) and PNUTS RNAi cells – and + Tet by DAPI. (B) Cell cycle analysis using flow cytometry. TbPNUTS RNAi cells treated with and without Tetracyclin for 2 days were pulse labeled with BrdU to label replicating cells. BrdU incorporation was detected using anti-BrdU antibody. Cells were stained with Propidium Iodide (PI) and analyzed by flow cytometry. (C) Cell cycle profile with PI channel only.

**S10 Fig. Ablation of TbPNUTS results in de-repression of silent MES and BES VSGs.** qRT-PCR analysis of MES VSG 1954, MES VSG 559, BES VSG MITat 1.1 and BES VSG MiTat 1.8 expression upon TbPNUTS ablation. Error bars indicate standard deviation from at least three experiments. P values were calculated using Student’s t test. ***, p value ≤ 0.001.

**S11 Fig. TbPNUTS inhibits transcription of VSG BES.** RNA-seq reads from the PNUTS RNAi were aligned to the *T. brucei* 427 BES sequences (14 BESs). Fold changes comparing plus and minus Tetracyclin were plotted over each BES as described in Fig 6C.

**S1 Table. Replicate JGT purification and MS analysis in *L. tarentolae*.** LtJGT was purified and proteins identified by mass spectrometry as in Table 1.

**S2 Table. *T. brucei* gene expression changes following PNUTS loss.** DEseq analysis comparing gene expression levels of the TbPNUTS RNAi cell line at day two plus and minus Tetracyclin. mRNAs that are at least 3-fold upregulated are listed, along with available gene descriptions and P values determined by Cuffdiff. No genes are downregulated.

## References

1. Shandilya J, Roberts SG. The transcription cycle in eukaryotes: from productive initiation to RNA polymerase II recycling. Biochim Biophys Acta. 2012;1819(5):391–400.

2. Kecman T, Kus K, Heo DH, Duckett K, Birot A, Liberatori S, et al. Elongation/Termination Factor Exchange Mediated by PP1 Phosphatase Orchestrates Transcription Termination. Cell Rep. 2018;25(1):259–69 e5.

3. Schreieck A, Easter AD, Etzold S, Wiederhold K, Lidschreiber M, Cramer P, et al. RNA polymerase II termination involves C-terminal-domain tyrosine dephosphorylation by CPF subunit Glc7. Nat Struct Mol Biol. 2014;21(2):175–9.

4. Bollen M, Peti W, Ragusa MJ, Beullens M. The extended PP1 toolkit: designed to create specificity. Trends in biochemical sciences. 2010;35(8):450–8.

5. Dancheck B, Nairn AC, Peti W. Detailed structural characterization of unbound protein phosphatase 1 inhibitors. Biochemistry. 2008;47(47):12346–56.

6. Ragusa MJ, Dancheck B, Critton DA, Nairn AC, Page R, Peti W. Spinophilin directs protein phosphatase 1 specificity by blocking substrate binding sites. Nat Struct Mol Biol. 2010;17(4):459–64.

7. Jagiello I, Beullens M, Stalmans W, Bollen M. Subunit structure and regulation of protein phosphatase-1 in rat liver nuclei. J Biol Chem. 1995;270(29):17257–63.

8. Kreivi JP, Trinkle-Mulcahy L, Lyon CE, Morrice NA, Cohen P, Lamond AI. Purification and characterisation of p99, a nuclear modulator of protein phosphatase 1 activity. FEBS Lett. 1997;420(1):57–62.

9. Lee JH, You J, Dobrota E, Skalnik DG. Identification and characterization of a novel human PP1 phosphatase complex. J Biol Chem. 2010;285(32):24466–76.

10. Lee JH, Skalnik DG. Wdr82 is a C-terminal domain-binding protein that recruits the Setd1A Histone H3-Lys4 methyltransferase complex to transcription start sites of transcribed human genes. Mol Cell Biol. 2008;28(2):609–18.

11. Dichtl B, Blank D, Ohnacker M, Friedlein A, Roeder D, Langen H, et al. A role for SSU72 in balancing RNA polymerase II transcription elongation and termination. Mol Cell. 2002;10(5):1139–50.

12. He X, Khan AU, Cheng H, Pappas DL, Jr., Hampsey M, Moore CL. Functional interactions between the transcription and mRNA 3’ end processing machineries mediated by Ssu72 and Sub1. Genes Dev. 2003;17(8):1030–42.

13. Vanoosthuyse V, Legros P, van der Sar SJ, Yvert G, Toda K, Le Bihan T, et al. CPF-associated phosphatase activity opposes condensin-mediated chromosome condensation. PLoS genetics. 2014;10(6):e1004415.

14. Nedea E, He X, Kim M, Pootoolal J, Zhong G, Canadien V, et al. Organization and function of APT, a subcomplex of the yeast cleavage and polyadenylation factor involved in the formation of mRNA and small nucleolar RNA 3’-ends. J Biol Chem. 2003;278(35):33000–10.

15. Cheng H, He X, Moore C. The essential WD repeat protein Swd2 has dual functions in RNA polymerase II transcription termination and lysine 4 methylation of histone H3. Mol Cell Biol. 2004;24(7):2932–43.

16. Nedea E, Nalbant D, Xia D, Theoharis NT, Suter B, Richardson CJ, et al. The Glc7 phosphatase subunit of the cleavage and polyadenylation factor is essential for transcription termination on snoRNA genes. Mol Cell. 2008;29(5):577–87.

17. Austenaa LM, Barozzi I, Simonatto M, Masella S, Della Chiara G, Ghisletti S, et al. Transcription of Mammalian cis-Regulatory Elements Is Restrained by Actively Enforced Early Termination. Mol Cell. 2015;60(3):460–74.

18. Campbell DA, Thomas S, Sturm NR. Transcription in kinetoplastid protozoa: why be normal? Microbes Infect. 2003;5(13):1231–40.

19. Borst P, Sabatini R. Base J: discovery, biosynthesis, and possible functions. Annu Rev Microbiol. 2008;62:235–51.

20. Cliffe LJ, Siegel TN, Marshall M, Cross GA, Sabatini R. Two thymidine hydroxylases differentially regulate the formation of glucosylated DNA at regions flanking polymerase II polycistronic transcription units throughout the genome of Trypanosoma brucei. Nucleic Acids Res. 2010;38(12):3923–35.

21. Reynolds D, Hofmeister BT, Cliffe L, Alabady M, Siegel TN, Schmitz RJ, et al. Histone H3 Variant Regulates RNA Polymerase II Transcription Termination and Dual Strand Transcription of siRNA Loci in Trypanosoma brucei. PLoS genetics. 2016;12(1):e1005758.

22. Siegel TN, Hekstra DR, Kemp LE, Figueiredo LM, Lowell JE, Fenyo D, et al. Four histone variants mark the boundaries of polycistronic transcription units in Trypanosoma brucei. Genes Dev. 2009;23(9):1063–76.

23. van Luenen HG, Farris C, Jan S, Genest PA, Tripathi P, Velds A, et al. Glucosylated hydroxymethyluracil, DNA base J, prevents transcriptional readthrough in Leishmania. Cell. 2012;150(5):909–21.

24. Reynolds D, Cliffe L, Forstner KU, Hon CC, Siegel TN, Sabatini R. Regulation of transcription termination by glucosylated hydroxymethyluracil, base J, in Leishmania major and Trypanosoma brucei. Nucleic Acids Res. 2014;42(15):9717–29.

25. Reynolds DL, Hofmeister BT, Cliffe L, Siegel TN, Anderson BA, Beverley SM, et al. Base J represses genes at the end of polycistronic gene clusters in Leishmania major by promoting RNAP II termination. Mol Microbiol. 2016;101(4):559–74.

26. Schulz D, Zaringhalam M, Papavasiliou FN, Kim HS. Base J and H3.V Regulate Transcriptional Termination in Trypanosoma brucei. PLoS genetics. 2016;12(1):e1005762.

27. Cliffe LJ, Hirsch G, Wang J, Ekanayake D, Bullard W, Hu M, et al. JBP1 and JBP2 Proteins Are Fe2+/2-Oxoglutarate-dependent Dioxygenases Regulating Hydroxylation of Thymidine Residues in Trypanosome DNA. J Biol Chem. 2012;287(24):19886–95.

28. Bullard W, da Rosa-Spiegler JL, Liu S, Wang D, Sabatini R. Identification of the glucosyltransferase that converts hydroxymethyluracil to base J in the trypanosomatid genome. JBC. 2014.

29. Sekar A, Merritt C, Baugh L, Stuart K, Myler PJ. Tb927.10.6900 encodes the glucosyltransferase involved in synthesis of base J in Trypanosoma brucei. Mol Biochem Parasitol. 2014;196(1):9–11.

30. Sabatini R, Cliffe L, Vainio L, Borst P. Enzymatic Formation of the Hypermodified DNA Base J. In: Grosjean H, editor. DNA and RNA Modification Enzymes: Comparative Structure, Mechanism, Function, Cellular Interactions and Evolution. Texas: Landes Biosciences; 2009. p. 120–31.

31. Cliffe LJ, Kieft R, Southern T, Birkeland SR, Marshall M, Sweeney K, et al. JBP1 and JBP2 are two distinct thymidine hydroxylases involved in J biosynthesis in genomic DNA of African trypanosomes. Nucleic Acids Res. 2009.

32. Kieft R, Brand V, Ekanayake DK, Sweeney K, DiPaolo C, Reznikoff WS, et al. JBP2, a SWI2/SNF2-like protein, regulates de novo telomeric DNA glycosylation in bloodstream form Trypanosoma brucei. Mol Biochem Parasitol. 2007;156(1):24–31.

33. Vainio S, Genest PA, ter Riet B, van Luenen H, Borst P. Evidence that J-binding protein 2 is a thymidine hydroxylase catalyzing the first step in the biosynthesis of DNA base J. Mol Biochem Parasitol. 2009;164(2):157–61.

34. Yu Z, Genest PA, ter Riet B, Sweeney K, DiPaolo C, Kieft R, et al. The protein that binds to DNA base J in trypanosomatids has features of a thymidine hydroxylase. Nucleic Acids Res. 2007;35(7):2107–15.

35. DiPaolo C, Kieft R, Cross M, Sabatini R. Regulation of trypanosome DNA glycosylation by a SWI2/SNF2-like protein. Mol Cell. 2005;17(3):441–51.

36. Heidebrecht T, Christodoulou E, Chalmers MJ, Jan S, Ter Riet B, Grover RK, et al. The structural basis for recognition of base J containing DNA by a novel DNA binding domain in JBP1. Nucleic Acids Res. 2011;39(13):5715–28.

37. Sabatini R, Meeuwenoord N, van Boom JH, Borst P. Recognition of base J in duplex DNA by J-binding protein. J Biol Chem. 2002;277:958–66.

38. Sabatini R, Meeuwenoord N, van Boom JH, Borst P. Site-specific interactions of JBP with base and sugar moieties in duplex J-DNA. Journal of Biological Chemistry. 2002;277:28150–6.

39. Bullard W, Cliffe L, Wang P, Wang Y, Sabatini R. Base J glucosyltransferase does not regulate the sequence specificity of J synthesis in trypanosomatid telomeric DNA. Mol Biochem Parasitol. 2015;204(2):77–80.

40. Bullard W, Kieft R, Sabatini R. A method for the efficient and selective identification of 5-hydroxymethyluracil in genomic DNA. Biol Methods Protoc. 2017;2(1).

41. van Leeuwen F, Kieft R, Cross M, Borst P. Biosynthesis and function of the modified DNA base beta-D-glucosyl-hydroxymethyluracil in Trypanosoma brucei. Molecular & Cellular Biology. 1998;18(10):5643–51.

42. Aphasizhev R, Aphasizheva I, Nelson RE, Gao G, Simpson AM, Kang X, et al. Isolation of a U-insertion/deletion editing complex from Leishmania tarentolae mitochondria. The EMBO Journal. 2003;22(4):913–24.

43. Hurley TD, Yang J, Zhang L, Goodwin KD, Zou Q, Cortese M, et al. Structural basis for regulation of protein phosphatase 1 by inhibitor-2. J Biol Chem. 2007;282(39):28874–83.

44. Terrak M, Kerff F, Langsetmo K, Tao T, Dominguez R. Structural basis of protein phosphatase 1 regulation. Nature. 2004;429(6993):780–4.

45. Jones DT, Cozzetto D. DISOPRED3: precise disordered region predictions with annotated protein-binding activity. Bioinformatics. 2015;31(6):857–63.

46. Ward JJ, McGuffin LJ, Bryson K, Buxton BF, Jones DT. The DISOPRED server for the prediction of protein disorder. Bioinformatics. 2004;20(13):2138–9.

47. Vacic V, Uversky VN, Dunker AK, Lonardi S. Composition Profiler: a tool for discovery and visualization of amino acid composition differences. BMC Bioinformatics. 2007;8:211.

48. Roy A, Kucukural A, Zhang Y. I-TASSER: a unified platform for automated protein structure and function prediction. Nat Protoc. 2010;5(4):725–38.

49. Yang J, Yan R, Roy A, Xu D, Poisson J, Zhang Y. The I-TASSER Suite: protein structure and function prediction. Nat Methods. 2015;12(1):7–8.

50. Brenchley R, Tariq H, McElhinney H, Szoor B, Huxley-Jones J, Stevens R, et al. The TriTryp phosphatome: analysis of the protein phosphatase catalytic domains. BMC Genomics. 2007;8:434.

51. Li Z, Tu X, Wang CC. Okadaic acid overcomes the blocked cell cycle caused by depleting Cdc2-related kinases in Trypanosoma brucei. Exp Cell Res. 2006;312(18):3504–16.

52. Schimanski B, Nguyen TN, Gunzl A. Highly efficient tandem affinity purification of trypanosome protein complexes based on a novel epitope combination. Eukaryot Cell. 2005;4(11):1942–50.

53. Ciurciu A, Duncalf L, Jonchere V, Lansdale N, Vasieva O, Glenday P, et al. PNUTS/PP1 regulates RNAPII-mediated gene expression and is necessary for developmental growth. PLoS genetics. 2013;9(10):e1003885.

54. Jerebtsova M, Klotchenko SA, Artamonova TO, Ammosova T, Washington K, Egorov VV, et al. Mass spectrometry and biochemical analysis of RNA polymerase II: targeting by protein phosphatase-1. Mol Cell Biochem. 2011;347(1-2):79–87.

55. Goos C, Dejung M, Janzen CJ, Butter F, Kramer S. The nuclear proteome of Trypanosoma brucei. PLoS One. 2017;12(7):e0181884.

56. Beullens M, Van Eynde A, Vulsteke V, Connor J, Shenolikar S, Stalmans W, et al. Molecular determinants of nuclear protein phosphatase-1 regulation by NIPP-1. J Biol Chem. 1999;274(20):14053–61.

57. Kim YM, Watanabe T, Allen PB, Kim YM, Lee SJ, Greengard P, et al. PNUTS, a protein phosphatase 1 (PP1) nuclear targeting subunit. Characterization of its PP1- and RNA-binding domains and regulation by phosphorylation. J Biol Chem. 2003;278(16):13819–28.

58. Liu J, Brautigan DL. Glycogen synthase association with the striated muscle glycogen-targeting subunit of protein phosphatase-1. Synthase activation involves scaffolding regulated by beta-adrenergic signaling. J Biol Chem. 2000;275(34):26074–81.

59. McAvoy T, Allen PB, Obaishi H, Nakanishi H, Takai Y, Greengard P, et al. Regulation of neurabin I interaction with protein phosphatase 1 by phosphorylation. Biochemistry. 1999;38(39):12943–9.

60. Huang G, Ulrich PN, Storey M, Johnson D, Tischer J, Tovar JA, et al. Proteomic analysis of the acidocalcisome, an organelle conserved from bacteria to human cells. PLoS Pathog. 2014;10(12):e1004555.

61. Mayer A, di Iulio J, Maleri S, Eser U, Vierstra J, Reynolds A, et al. Native elongating transcript sequencing reveals human transcriptional activity at nucleotide resolution. Cell. 2015;161(3):541–54.

62. Neil H, Malabat C, d’Aubenton-Carafa Y, Xu Z, Steinmetz LM, Jacquier A. Widespread bidirectional promoters are the major source of cryptic transcripts in yeast. Nature. 2009;457(7232):1038–42.

63. Nojima T, Gomes T, Grosso ARF, Kimura H, Dye MJ, Dhir S, et al. Mammalian NET-Seq Reveals Genome-wide Nascent Transcription Coupled to RNA Processing. Cell. 2015;161(3):526–40.

64. Preker P, Nielsen J, Kammler S, Lykke-Andersen S, Christensen MS, Mapendano CK, et al. RNA exosome depletion reveals transcription upstream of active human promoters. Science. 2008;322(5909):1851–4.

65. Xu Z, Wei W, Gagneur J, Perocchi F, Clauder-Munster S, Camblong J, et al. Bidirectional promoters generate pervasive transcription in yeast. Nature. 2009;457(7232):1033–7.

66. Kolev NG, Franklin JB, Carmi S, Shi H, Michaeli S, Tschudi C. The transcriptome of the human pathogen Trypanosoma brucei at single-nucleotide resolution. PLoS Pathog. 2010;6(9).

67. Wedel C, Forstner KU, Derr R, Siegel TN. GT-rich promoters can drive RNA pol II transcription and deposition of H2A.Z in African trypanosomes. EMBO J. 2017;36(17):2581–94.

68. Gros J, Kumar C, Lynch G, Yadav T, Whitehouse I, Remus D. Post-licensing Specification of Eukaryotic Replication Origins by Facilitated Mcm2-7 Sliding along DNA. Mol Cell. 2015;60(5):797–807.

69. Mori S, Shirahige K. Perturbation of the activity of replication origin by meiosis-specific transcription. J Biol Chem. 2007;282(7):4447–52.

70. Nieduszynski CA, Blow JJ, Donaldson AD. The requirement of yeast replication origins for pre-replication complex proteins is modulated by transcription. Nucleic Acids Res. 2005;33(8):2410–20.

71. Nieduszynski CA, Knox Y, Donaldson AD. Genome-wide identification of replication origins in yeast by comparative genomics. Genes Dev. 2006;20(14):1874–9.

72. Snyder M, Sapolsky RJ, Davis RW. Transcription interferes with elements important for chromosome maintenance in Saccharomyces cerevisiae. Mol Cell Biol. 1988;8(5):2184–94.

73. Tiengwe C, Marcello L, Farr H, Dickens N, Kelly S, Swiderski M, et al. Genome-wide analysis reveals extensive functional interaction between DNA replication initiation and transcription in the genome of Trypanosoma brucei. Cell Rep. 2012;2(1):185–97.

74. Lombrana R, Alvarez A, Fernandez-Justel JM, Almeida R, Poza-Carrion C, Gomes F, et al. Transcriptionally Driven DNA Replication Program of the Human Parasite Leishmania major. Cell Rep. 2016;16(6):1774–86.

75. Horn D. Antigenic variation in African trypanosomes. Mol Biochem Parasitol. 2014;195(2):123–9.

76. Horn D, McCulloch R. Molecular mechanisms underlying the control of antigenic variation in African trypanosomes. Curr Opin Microbiol. 2010;13(6):700–5.

77. Muller LSM, Cosentino RO, Forstner KU, Guizetti J, Wedel C, Kaplan N, et al. Genome organization and DNA accessibility control antigenic variation in trypanosomes. Nature. 2018;563(7729):121–5.

78. Cross GA, Kim HS, Wickstead B. Capturing the variant surface glycoprotein repertoire (the VSGnome) of Trypanosoma brucei Lister 427. Mol Biochem Parasitol. 2014;195(1):59–73.

79. Wickstead B, Ersfeld K, Gull K. The small chromosomes of Trypanosoma brucei involved in antigenic variation are constructed around repetitive palindromes. Genome Res. 2004;14(6):1014–24.

80. Hertz-Fowler C, Figueiredo LM, Quail MA, Becker M, Jackson A, Bason N, et al. Telomeric expression sites are highly conserved in Trypanosoma brucei. PLoS One. 2008;3(10):e3527.

81. Ehlers B, Czichos J, Overath P. RNA turnover in Trypanosoma brucei. Mol Cell Biol. 1987;7(3):1242–9.

82. Kabiri M, Steverding D. Studies on the recycling of the transferrin receptor in Trypanosoma brucei using an inducible gene expression system. Eur J Biochem. 2000;267(11):3309–14.

83. Kerry LE, Pegg EE, Cameron DP, Budzak J, Poortinga G, Hannan KM, et al. Selective inhibition of RNA polymerase I transcription as a potential approach to treat African trypanosomiasis. PLoS Negl Trop Dis. 2017;11(3):e0005432.

84. Vanhamme L, Berberof M, Le Ray D, Pays E. Stimuli of differentiation regulate RNA elongation in the transcription units for the major stage-specific antigens of Trypanosoma brucei. Nucleic Acids Res. 1995;23(11):1862–9.

85. Vanhamme L, Pays E, McCulloch R, Barry JD. An update on antigenic variation in African trypanosomes. TRENDS in Parasitology. 2001;17:338–43.

86. Vanhamme L, Poelvoorde P, Pays A, Tebabi P, Van Xong H, Pays E. Differential RNA elongation controls the variant surface glycoprotein gene expression sites of Trypanosoma brucei. Mol Microbiol. 2000;36(2):328–40.

87. Clayton C, Shapira M. Post-transcriptional regulation of gene expression in trypanosomes and leishmanias. Mol Biochem Parasitol. 2007;156(2):93–101.

88. Ruthenburg AJ, Wang W, Graybosch DM, Li H, Allis CD, Patel DJ, et al. Histone H3 recognition and presentation by the WDR5 module of the MLL1 complex. Nat Struct Mol Biol. 2006;13(8):704–12.

89. Soares LM, Buratowski S. Yeast Swd2 is essential because of antagonism between Set1 histone methyltransferase complex and APT (associated with Pta1) termination factor. J Biol Chem. 2012;287(19):15219–31.

90. Ng HH, Robert F, Young RA, Struhl K. Targeted recruitment of Set1 histone methylase by elongating Pol II provides a localized mark and memory of recent transcriptional activity. Mol Cell. 2003;11(3):709–19.

91. Dingar D, Tu WB, Resetca D, Lourenco C, Tamachi A, De Melo J, et al. MYC dephosphorylation by the PP1/PNUTS phosphatase complex regulates chromatin binding and protein stability. Nature communications. 2018;9(1):3502.

92. Parua PK, Booth GT, Sanso M, Benjamin B, Tanny JC, Lis JT, et al. A Cdk9-PP1 switch regulates the elongation-termination transition of RNA polymerase II. Nature. 2018;558(7710):460–4.

93. Srivastava A, Badjatia N, Lee JH, Hao B, Gunzl A. An RNA polymerase II-associated TFIIF-like complex is indispensable for SL RNA gene transcription in Trypanosoma brucei. Nucleic Acids Res. 2017.

94. Urbaniak MD, Martin DM, Ferguson MA. Global quantitative SILAC phosphoproteomics reveals differential phosphorylation is widespread between the procyclic and bloodstream form lifecycle stages of Trypanosoma brucei. J Proteome Res. 2013;12(5):2233–44.

95. Guo Z, Stiller JW. Comparative genomics of cyclin-dependent kinases suggest co-evolution of the RNAP II C-terminal domain and CTD-directed CDKs. BMC Genomics. 2004;5:69.

96. Das A, Banday M, Fisher MA, Chang YJ, Rosenfeld J, Bellofatto V. An essential domain of an early-diverged RNA polymerase II functions to accurately decode a primitive chromatin landscape. Nucleic Acids Res. 2017;45(13):7886–96.

97. Das A, Bellofatto V. The non-canonical CTD of RNAP-II is essential for productive RNA synthesis in Trypanosoma brucei. PLoS One. 2009;4(9):e6959.

98. Rocha AA, Moretti NS, Schenkman S. Stress induces changes in the phosphorylation of Trypanosoma cruzi RNA polymerase II, affecting its association with chromatin and RNA processing. Eukaryot Cell. 2014;13(7):855–65.

99. Core LJ, Waterfall JJ, Lis JT. Nascent RNA sequencing reveals widespread pausing and divergent initiation at human promoters. Science. 2008;322(5909):1845–8.

100. Preker P, Nielsen J, Schierup MH, Jensen TH. RNA polymerase plays both sides: vivid and bidirectional transcription around and upstream of active promoters. Cell Cycle. 2009;8(8):1106–7.

101. Sigova AA, Mullen AC, Molinie B, Gupta S, Orlando DA, Guenther MG, et al. Divergent transcription of long noncoding RNA/mRNA gene pairs in embryonic stem cells. Proc Natl Acad Sci U S A. 2013;110(8):2876–81.

102. Wei W, Pelechano V, Jarvelin AI, Steinmetz LM. Functional consequences of bidirectional promoters. Trends Genet. 2011;27(7):267–76.

103. Flynn RA, Almada AE, Zamudio JR, Sharp PA. Antisense RNA polymerase II divergent transcripts are P-TEFb dependent and substrates for the RNA exosome. Proc Natl Acad Sci U S A. 2011;108(26):10460–5.

104. Ntini E, Jarvelin AI, Bornholdt J, Chen Y, Boyd M, Jorgensen M, et al. Polyadenylation site-induced decay of upstream transcripts enforces promoter directionality. Nat Struct Mol Biol. 2013;20(8):923–8.

105. Schulz D, Schwalb B, Kiesel A, Baejen C, Torkler P, Gagneur J, et al. Transcriptome surveillance by selective termination of noncoding RNA synthesis. Cell. 2013;155(5):1075–87.

106. Descostes N, Heidemann M, Spinelli L, Schuller R, Maqbool MA, Fenouil R, et al. Tyrosine phosphorylation of RNA polymerase II CTD is associated with antisense promoter transcription and active enhancers in mammalian cells. Elife. 2014;3:e02105.

107. Fong N, Saldi T, Sheridan RM, Cortazar MA, Bentley DL. RNA Pol II Dynamics Modulate Co-transcriptional Chromatin Modification, CTD Phosphorylation, and Transcriptional Direction. Mol Cell. 2017;66(4):546–57 e3.

108. Shah N, Maqbool MA, Yahia Y, El Aabidine AZ, Esnault C, Forne I, et al. Tyrosine-1 of RNA Polymerase II CTD Controls Global Termination of Gene Transcription in Mammals. Mol Cell. 2018;69(1):48–61 e6.

109. Shetty A, Kallgren SP, Demel C, Maier KC, Spatt D, Alver BH, et al. Spt5 Plays Vital Roles in the Control of Sense and Antisense Transcription Elongation. Mol Cell. 2017;66(1):77–88 e5.

110. Dellino GI, Cittaro D, Piccioni R, Luzi L, Banfi S, Segalla S, et al. Genome-wide mapping of human DNA-replication origins: levels of transcription at ORC1 sites regulate origin selection and replication timing. Genome Res. 2013;23(1):1–11.

111. Miotto B, Ji Z, Struhl K. Selectivity of ORC binding sites and the relation to replication timing, fragile sites, and deletions in cancers. Proc Natl Acad Sci U S A. 2016;113(33):E4810–9.

112. Soudet J, Gill JK, Stutz F. Noncoding transcription influences the replication initiation program through chromatin regulation. Genome Res. 2018;28(12):1882–93.

113. Kim HS. Genome-wide function of MCM-BP in Trypanosoma brucei DNA replication and transcription. Nucleic Acids Res. 2019;47(2):634–47.

114. Benmerzouga I, Concepcion-Acevedo J, Kim HS, Vandoros AV, Cross GA, Klingbeil MM, et al. Trypanosoma brucei Orc1 is essential for nuclear DNA replication and affects both VSG silencing and VSG switching. Mol Microbiol. 2013;87(1):196–210.

115. Anderson SJ, Sikes ML, Zhang Y, French SL, Salgia S, Beyer AL, et al. The transcription elongation factor Spt5 influences transcription by RNA polymerase I positively and negatively. J Biol Chem. 2011;286(21):18816–24.

116. Viktorovskaya OV, Appling FD, Schneider DA. Yeast transcription elongation factor Spt5 associates with RNA polymerase I and RNA polymerase II directly. J Biol Chem. 2011;286(21):18825–33.

117. Wirtz E, Leal S, Ochatt C, Cross GA. A tightly regulated inducible expression system for conditional gene knock-outs and dominant-negative genetics in Trypanosoma brucei. Mol Biochem Parasitol. 1999;99(1):89–101.

118. Wickstead B, Ersfeld K, Gull K. Targeting of a tetracycline-inducible expression system to the transcriptionally silent minichromosomes of Trypanosoma brucei. Mol Biochem Parasitol. 2002;125(1-2):211–6.

119. Oberholzer M, Morand S, Kunz S, Seebeck T. A vector series for rapid PCR-mediated C-terminal in situ tagging of Trypanosoma brucei genes. Mol Biochem Parasitol. 2006;145(1):117–20.

120. Lim JM, Sherling D, Teo CF, Hausman DB, Lin D, Wells L. Defining the regulated secreted proteome of rodent adipocytes upon the induction of insulin resistance. J Proteome Res. 2008;7(3):1251–63.

121. Baez-Santos YM, Mielech AM, Deng X, Baker S, Mesecar AD. Catalytic function and substrate specificity of the papain-like protease domain of nsp3 from the Middle East respiratory syndrome coronavirus. J Virol. 2014;88(21):12511–27.

122. Chen S, Zhou Y, Chen Y, Gu J. fastp: an ultra-fast all-in-one FASTQ preprocessor. Bioinformatics. 2018;34(17):i884–i90.

123. Langmead B, Salzberg SL. Fast gapped-read alignment with Bowtie 2. Nature methods. 2012;9(4):357–9.

124. Li H, Handsaker B, Wysoker A, Fennell T, Ruan J, Homer N, et al. The Sequence Alignment/Map format and SAMtools. Bioinformatics. 2009;25(16):2078–9.

125. Ryba T, Battaglia D, Pope BD, Hiratani I, Gilbert DM. Genome-scale analysis of replication timing: from bench to bioinformatics. Nat Protoc. 2011;6(6):870–95.

